# Deep mutational analysis reveals functional trade-offs in the sequences of EGFR autophosphorylation sites

**DOI:** 10.1101/273516

**Authors:** Aaron J. Cantor, Neel H. Shah, John Kuriyan

## Abstract

Upon activation, the epidermal growth factor receptor (EGFR) phosphorylates tyrosine residues in its cytoplasmic tail, which triggers the binding of Src Homology 2 (SH2) and Phosphotyrosine Binding (PTB) domains and initiates downstream signaling. The sequences flanking the tyrosine residues (referred to as phosphosites) must be compatible with phosphorylation by the EGFR kinase domain and the recruitment of adapter proteins, while minimizing phosphorylation that would reduce the fidelity of signal transmission. In order to understand how phosphosite sequences encode these functions within a small set of residues, we carried out high-throughput mutational analysis of three phosphosite sequences in the EGFR tail. We used bacterial surface-display of peptides, coupled with deep sequencing, to monitor phosphorylation efficiency and the binding of the SH2 and PTB domains of the adapter proteins Grb2 and Shc1, respectively. We found that the sequences of phosphosites in the EGFR tail are restricted to a subset of the range of sequences that can be phosphorylated efficiently by EGFR. Although efficient phosphorylation by EGFR can occur with either acidic or large hydrophobic residues at the −1 position with respect to the tyrosine, hydrophobic residues are generally excluded from this position in tail sequences. The mutational data suggest that this restriction results in weaker binding to adapter proteins, but also disfavors phosphorylation by the cytoplasmic tyrosine kinases c-Src and c-Abl. Our results show how EGFR-family phosphosites achieve a trade-off between minimizing off-pathway phosphorylation while maintaining the ability to recruit the diverse complement of effectors required for downstream pathway activation.

## Introduction

The epidermal growth factor receptor (EGFR) is a tyrosine kinase that couples extracellular ligand binding to activation of intracellular signaling pathways (1–3). Activation of human EGFR, also called ErbB1 or Her1 (Human EGF Receptor 1), results from ligand-induced homodimerization or heterodimerization with one of three family members, Her2/ErbB2/neu, Her3/ErbB3, or Her4/ErbB4 (4). Allosteric activation of one kinase domain within this dimer by the other kinase domain results in autophosphorylation of multiple tyrosines within the C-terminal tails of both monomers (5). The resulting phosphotyrosine and flanking residues (phosphosites) can then serve as binding sites for intracellular proteins containing Src Homology 2 (SH2) or Phosphotyrosine Binding (PTB) domains (6, 7). With their enzymatic or scaffolding activities recruited to the plasma membrane, these effector proteins propagate signals inside the cell (8) (**Figure 1A**).

**Figure 1:**
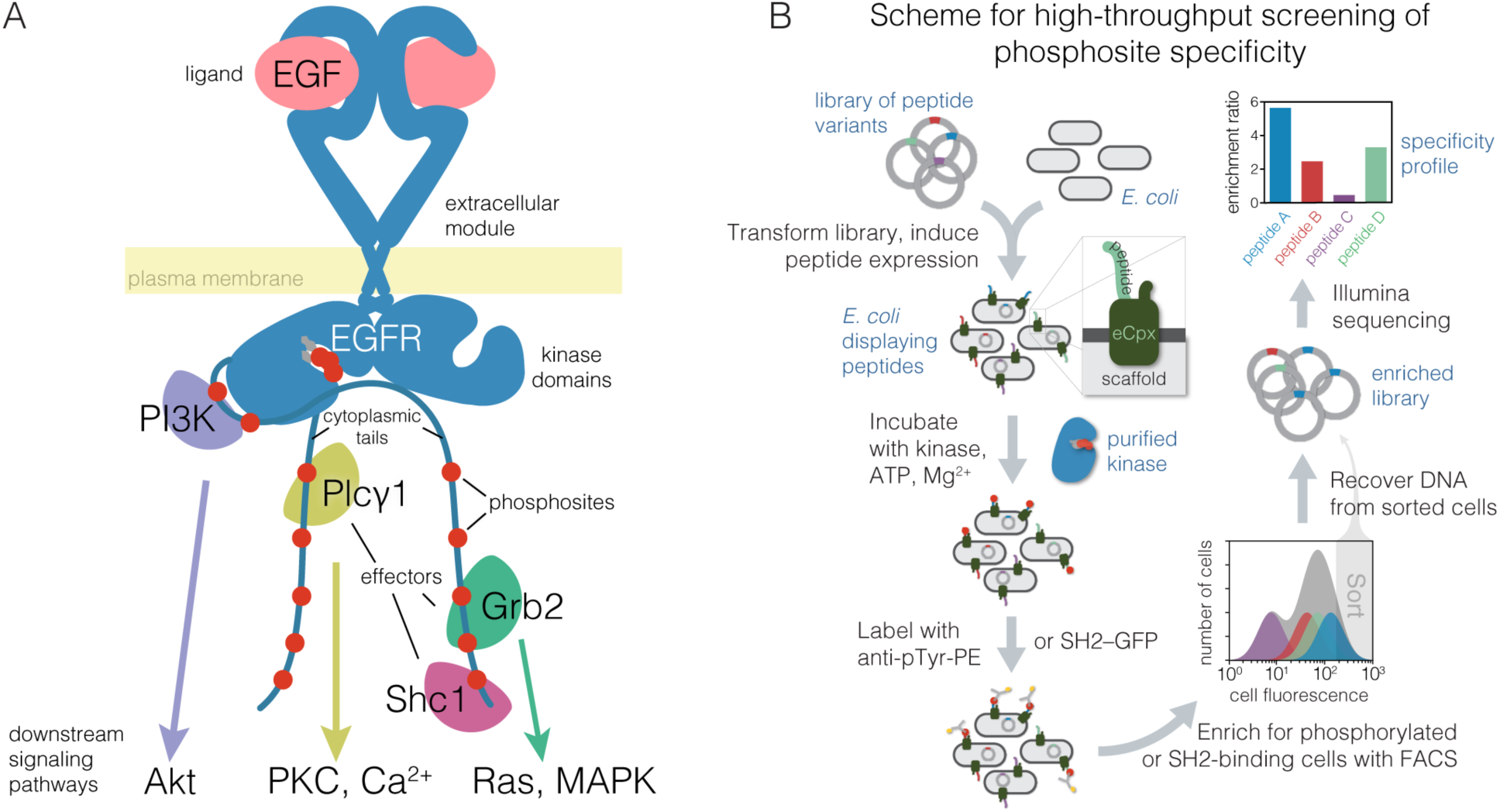
Overview of EGFR signal transduction at the membrane and a bacterial surface-display scheme to analyze specificity of tyrosine kinases and phosphotyrosine binding proteins. A. Illustration of membrane-proximal EGFR signaling components. Autophosphorylation of the tyrosine phosphosites in the C-terminal cytoplasmic tail (red circles) by the activated kinase domain produces binding sites for many downstream effectors, a subset of which are depicted. These effectors go on to activate second-messenger pathways, also depicted. PI3K, phosphoinositide 3-kinase regulatory subunit; Plcγ1, phospholipase C-gamma-1; Grb2,growth factor receptor-bound protein 2; Shc1, Src homology 2 domain-containing-transforming protein C1; PKC, protein kinase C; MAPK, mitogen activated protein kinases. **B.** Workflow for determining phosphosite specificity profiles of tyrosine kinases and phosphotyrosine binding proteins by bacterial surface-display coupled with fluorescence activated cell sorting and deep sequencing. Phosphotyrosine on the surface of the cells is detected either by immunostaining with an anti-phosphotyrosine antibody, or, for binding profiles, with a tandem SH2– or PTB–GFP construct. The frequency of each peptide-coding sequence in the highly phosphorylated population, or enrichment, and thus the relative efficiency of phosphorylation or binding for each peptide, is determined by counting the number of sequencing reads for each peptide in the sorted and unsorted populations.

Tyrosine kinases recognize their substrates through the formation of a short antiparallel *β*-stranded interaction between the substrate peptide and the activation loop of the kinase domain (9). The kinase domain provides limited opportunity for stereospecific engagement of substrate sidechains. This fact contributes to the impression that tyrosine kinases are “sloppy enzymes,” (10, 11) and is consistent with the failure to develop high-affinity substrate-mimicking inhibitors. The catalytic domains of tyrosine kinases do have intrinsic preferences for some substrate sequences over others, with specificity being determined by the pattern of amino acid residues directly adjacent to the tyrosine (12, 13).

For EGFR family members, the high local concentration of tail phosphosites with respect to the kinase domains may allow these sites to be phosphorylated without much regard to sequence. This made us wonder about the extent to which each phosphosite in the cytoplasmic tails of EGFR-family members is optimized in its sequence for the recruitment of specific adaptor proteins versus phosphorylation efficiency by the EGFR kinase domain. The subset of tail tyrosines that are phosphorylated during EGFR signaling is thought to define the subset of downstream pathways that are activated, although the extent to which the intrinsic specificity of the kinase domain of EGFR allows discrimination between different phosphosites in the tail has not been mapped (14, 15). The properties that determine the efficiency of binding of SH2 or PTB domains to EGFR-family phosphosites are also not completely understood, because several SH2 or PTB domains can bind to a particular phosphosite, and each SH2 and PTB domain can use multiple different phosphosites (16–20).

Another layer of complexity in EGFR signaling is the potential for cross-talk with cytoplasmic tyrosine kinases, particularly the ubiquitously expressed kinases c-Src and c-Abl (21, 22). Direct phosphorylation of EGFR by c-Src might allow transactivation of EGFR in the absence of growth factors (23–25). Recently, c-Src has been shown to supply a priming phosphorylation that improves EGFR catalytic efficiency for a phosphosite with two neighboring tyrosine residues in the adapter protein Shc1 (26). Despite the evident importance of c-Src in EGFR signaling, very few phosphosites in EGFR family members have been shown to be direct substrates of c-Src (27–29). In particular, the inhibition of Src-family kinases has little effect on the phosphorylation of EGFR tail phosphosites (30). We therefore wondered whether there are mechanisms that insulate phosphosites in the EGFR-family receptors from phosphorylation by c-Src and other cytoplasmic tyrosine kinases.

In this work, we address several questions about the phosphosites in the cytoplasmic tails of EGFR-family receptors. What is the intrinsic specificity of the EGFR kinase domain with respect to all potential substrates? How does the intrinsic specificity of EGFR map onto tail phosphosites, and how is this specificity differentiated from that of c-Src and c-Abl? How has the specificity of SH2 and PTB domains impinged on the evolution of the sequences of EGFR-family tail phosphosites? To answer these questions, we assayed the activity of EGFR against thousands of defined sequences, representing tyrosine phosphorylation sites in the human proteome, using a high-throughput method based on bacterial surface-display of peptides and deep sequencing (**Figure 1B**). This method was used recently to determine the mechanistic basis for the orthogonal specificity of two kinases, Lck and ZAP-70, in the T-cell receptor pathway (31) and to map the specificity of kinases in the Src family (32). To delineate the specificities of SH2 and PTB domains, we augmented the bacterial surface-display system to measure protein binding instead of phosphorylation efficiency.

Deep mutational scanning of EGFR phosphosites, with selection based on either phosphorylation or protein binding, revealed specificity determinants that differ from optimal motifs observed in previous studies (12, 26, 33). We find that phosphosites in the tails of EGFR family members avoid sequence features that are consistent with efficient phosphorylation by EGFR and efficient binding of the Shc1 PTB domain and the Grb2 SH2 domain. The sequence features that are avoided in the tails would, if present, promote phosphorylation by c-Src. Thus, our studies of specificity in the context of natural phosphosite sequences have uncovered evidence of evolutionary trade-offs between on-pathway phosphorylation and binding, and suppression of potentially interfering reactivity in EGFR signaling.

## Results and Discussion

### Specificity profile of the EGFR kinase domain derived from a library of tyrosine phosphorylation sites found in the human proteome

Previous studies of tyrosine kinase specificity have been based primarily on degenerate peptide libraries, in which a tyrosine residue is flanked by random amino acid residues, except at one defined position (34). This approach assesses the sufficiency of particular types of residues at specific sites to confer phosphorylation by the kinase of interest. To gain an understanding of EGFR kinase specificity in the context of defined sequences, we developed a high-throughput assay, based on bacterial surface-display (35, 36), fluorescence-activated cell sorting, and deep sequencing, to screen for efficiently phosphorylated tyrosine phosphosites (31) (**Figure 1B**). This multiplexed bacterial surface-display kinase assay has been used recently to characterize tyrosine kinase specificity in the T-cell receptor signaling pathway (31), and to analyze tyrosine kinase specificity on substrates spanning the human proteome (32).

We used this method to screen the EGFR kinase against a library of 15-residue tyrosine-containing peptides, referred to as the “Human-pTyr library”. This library corresponds to a diverse set of ∼2600 tyrosine-containing sequences from the human proteome that have been reported as tyrosine kinase substrates in the PhosphoSitePlus (37) or Uniprot (38) databases ((32) and see Methods). Briefly, *E. coli* cells displaying individual peptides from the Human-pTyr library on their surfaces were subjected to phosphorylation by the purified EGFR kinase. The cells were then labeled with an anti-phosphotyrosine antibody, and the highly phosphorylated population was enriched by fluorescence-activated cell sorting. The abundance of each peptide-coding DNA sequence in the sorted and unsorted samples were inferred by their read frequencies in high-throughput sequencing. The ratio of sorted over input read frequency gives an enrichment score for each peptide. This score correlates well with *in vitro* measurement of kinase specific activity at low peptide concentrations relative to expected *K*M values across a wide dynamic range (**Supplemental Figure S1A**), indicating that it is a good measure of catalytic efficiency. The library contains ∼700 sequences with more than one tyrosine residue in a 15-residue stretch, and these were analyzed separately.

A key step in our analysis of EGFR specificity was the use of a soluble, dimeric form of the EGFR intracellular module. The isolated EGFR kinase domain has been shown to have ∼15 times higher specific activity when forced to dimerize on lipid vesicles, compared to the activity of the monomeric kinase domain (5, 39, 40). We designed constructs that consist of the EGFR intracellular module fused C-terminally to the proteins FKBP and FRB, which when mixed together along with rapamycin exhibit a ∼30-fold higher specific activity than either protein alone (**Supplemental Figure S1B**). This soluble construct also includes part of the juxtamembrane region, the kinase domain, and the full-length cytoplasmic tail (see **Supplemental Methods** for details). As discussed below, this dimerized construct of EGFR has comparable specific activity against preferred substrates to that of the kinase domain of c-Src against its preferred substrates.

The distribution of phosphorylation enrichment values for single-tyrosine peptides in the Human-pTyr library screened against human EGFR kinase is centered close to zero. The distribution has a long tail toward higher values, suggesting that EGFR phosphorylates most sites poorly, and phosphorylates a relatively narrow subset efficiently (**Figure 2A**). To determine the sequence features that underlie efficient phosphorylation by EGFR, we analyzed the positional enrichment of amino acid residues in peptides in the top quartile of enrichment ratios in two replicates (**Figure 2B**). The data are displayed using a sequence logo diagram (“phoshpo-pLogo”), which compares the frequency of an amino acid residue at each position in the set of efficiently-phosphorylated sequences, relative to the frequency of that residue in the same position with respect to tyrosine residues in a reference database of tyrosine-containing sequences (41). For clarity, we refer to logos generated from sequences that are filtered by experimental data on phosphorylation efficiency as “phospho-pLogos”, and those based on a collection of sequences derived from bioinformatics analysis as “sequence pLogos.”

**Figure 2:**
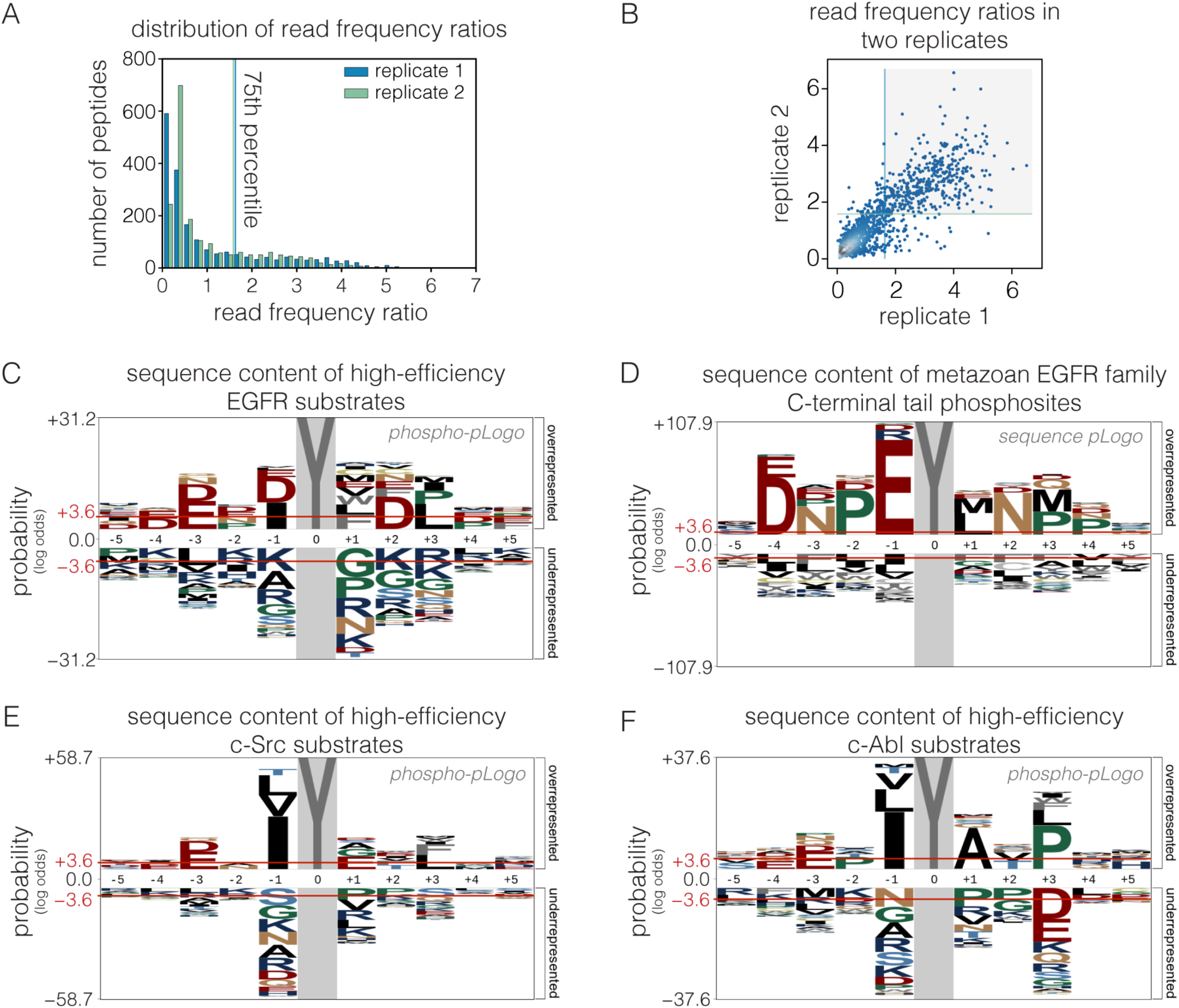
Comparison of intrinsic EGFR and c-Src substrate specificity with EGFR-family phosphosite sequences. A. Histogram of peptide read frequency ratios from EGFR phosphorylation of a library of human phosphosites obtained by bacterial surface-display and deep sequencing. The distribution of ratios of read frequencies for input and sorted samples are plotted from two replicate experiments. **B.** Read frequency ratios for two replicate Human-pTyr library phosphorylation experiments plotted against each other. Peptides with ratios above the 75^th^ percentile in both replicates (gray box) were counted as highly phosphorylated in C. **C.** Phosphorylation probability logo (“phospho-pLogo”) of highly phosphorylated peptides for EGFR in the bacterial surface-display experiment. The height of each letter corresponds to the negative log-odds ratio of binomial probabilities of finding a given amino acid residue at a particular sequence position at higher versus lower frequencies than the expected positional frequency for all peptides in the library. Higher values indicate an enrichment of a residue versus the background distribution. Red lines indicate the log-odds ratio values for a significance level of 0.05, as defined in (41). **D.** Sequence probability logo (“sequence pLogo”) of EGFR-family C-terminal tail tyrosines. Sequence segments surrounding tyrosine were extracted from the regions C-terminal to the kinase domain for metazoan EGFR-family protein sequences. The positional amino acid frequency in these segments was compared to the frequency in metazoan intracellular and transmembrane proteins and plotted as a pLogo. **E.** and **F.** Phospho-pLogo of highly phosphorylated sequences from c-Src phosphorylation (**E**) and c-Abl phosphorylation (**F**) of the pan-pY library (raw data from (32)). Sequences above the 75^th^ percentile in three replicates are included in the highly-phosphorylated set.

Tyrosine kinases have a general preference for peptides with negatively-charged residues located before the tyrosine (42). Consistent with this, acidic residues are enriched and basic residues are depleted in the phospho-pLogo diagram corresponding to the set of peptides that are phosphorylated efficiently by EGFR (**Figure 2C**). The phosphorylation motif determined for EGFR in earlier work using oriented peptide libraries (12), with acidic residues before the tyrosine and a hydrophobic residue in the position immediately after the tyrosine (the +1 position), is also apparent. In our data, peptides containing a −1 acidic/+1 hydrophobic motif are significantly more likely to be in the highly phosphorylated set (*p*<10**^−16^**, Fisher’s Exact Test), suggesting that this feature is predictive of efficient phosphorylation by EGFR. Unexpectedly, peptides with a −1 isoleucine or +3 leucine residue are also significantly enriched among sites that are phosphorylated efficiently by EGFR (*p*<10**^−14^**, Fisher’s Exact Test). Sequences with a leucine at the +3 position would be compatible with the binding of several SH2 domains, such as those of c-Src and phosphatidylinositol-3’-kinase (43), suggesting convergence in specificity between EGFR phosphorylation and SH2 domain binding in this instance.

When peptides with more than one tyrosine are included in the analysis, tyrosine is enriched significantly at multiple positions in the resulting phospho-pLogo (**Supplemental Figure S2).** This might reflect, in part, the recently described preference of EGFR for a Tyr–phospho-Tyr motif (26); we do not see an equivalent enrichment of tyrosine residues in the phospho-pLogo diagrams for other tyrosine kinases tested using the Human-pTyr library and bacterial surface-display (32). The analysis in this paper is restricted to the set of sequences that contain only one tyrosine residue, at the central position. This includes many phosphosites in which a secondary tyrosine is mutated to alanine in the library.

### Phosphosites in the cytoplasmic tails of EGFR-family members represent a subset of sequence patterns that are phosphorylated efficiently by EGFR

We next compared the sequence patterns of the highly phosphorylated peptides in the screen of human phosphosites to the sequence patterns of phosphosites in members of the EGFR family. We analyzed the positional amino acid enrichment in the 10 residues flanking each C-terminal tyrosine in a diverse collection of 87 EGFR family members from across metazoan evolution, displayed as a sequence-pLogo diagram in **Figure 2D**. The sequence-pLogo diagram for tail phosphosites is different from the phospho-pLogo for efficiently-phosphorylated EGFR substrates. Note, however, that the conserved features of the tail sequences comprise a subset of the sequence features that define efficient phosphorylation by the EGFR kinase domain. The major difference between the two logos arises from the enrichment of a central “EYL” motif in the phosphosites of EGFR-family tails. This EYL motif is compatible with the phospho-pLogo for EGFR (**Figure 2C**), as well as the optimal EEEEYFLVE motif reported for EGFR based on *in vitro* phosphorylation of degenerate peptide libraries (12, 44). The convergence of EGFR-family phosphosites on the EYL motif implies that efficient phosphorylation by EGFR is an important evolutionary pressure shaping these sequences.

Peptides with an isoleucine residue at the −1 position, rather than a glutamate, are also phosphorylated efficiently by EGFR, but a hydrophobic residue is very rarely found at the −1 position in the tails of EGFR-family members. The preferred motifs for phosphorylation by cytoplasmic tyrosine kinases often consist of hydrophobic residues at the −1 and +3 positions (Songyang et al. 1995; Miller et al. 2008; Deng et al. 2014). The phospho-pLogo diagrams for c-Src and c-Abl phosphorylation of the human-pTyr library confirm the importance of a large hydrophobic residue at the −1 position for efficient phosphorylation ((32), **Figure 2E, 2F**). c-Src and c-Abl also disfavor large hydrophobic residues at the +1 position (**Figure 2E, 2F**), which are overrepresented in the sequence-pLogo for EGFR-family tail phosphosites. These observations suggest that the EYL motif in EGFR-family tails may have arisen from selection pressure to minimize phosphorylation by cytoplasmic tyrosine kinases, such as c-Src and c-Abl.

Other differences between the phospho-pLogo for efficient EGFR phosphorylation and the sequence pLogo for EGFR-family tail phosphosites arise from the clear imprint of the binding motifs for SH2 and PTB domains in the tail phosphosites. For example, the binding motifs for the SH2 domains of Grb2 (pYxN, where x is any residue) and phosphatidylinositol-3’-kinases (pYxxF, where F is any hydrophobic residue) (45, 46), as well as for the PTB domains of Shc and Dok proteins (NPxpY) (7, 47), are highly enriched in the sequence-pLogo (**Figure 2D**). Because the sequence-pLogo is derived from EGFR-family sequences spread across the metazoan lineage, this result points to a conservation in the core specificity determinants of SH2 and PTB domains across evolution.

### Saturation mutagenesis of EGFR tail phosphosites

We performed mutational screens to assess the contribution to phosphorylation efficiency of each position in three phosphosites in the EGFR tail (Tyr 992, Tyr 1086, and Tyr 1114) (**Figure 3A**). Of these, the sequence flanking Tyr 992 is most similar to the phospho-pLogo derived from the Human-pTyr library (**Figure 2C**) and previously published motifs, with an EYL motif and mostly acidic residues upstream of the tyrosine (12, 26). The other two sites, spanning Tyr 1086 and Tyr 1114, both contain an NPxY motif, the signature of PTB domain binding, as well as a +2 asparagine, which is the main determinant for Grb2 SH2 binding. These two sites differ in their-1 and +1 residues, however. The Tyr 1114 phosphosite contains the consensus EYL motif, while Tyr 1086 diverges from this consensus, with valine and histidine at the −1 and +1 positions, respectively.

**Figure 3:**
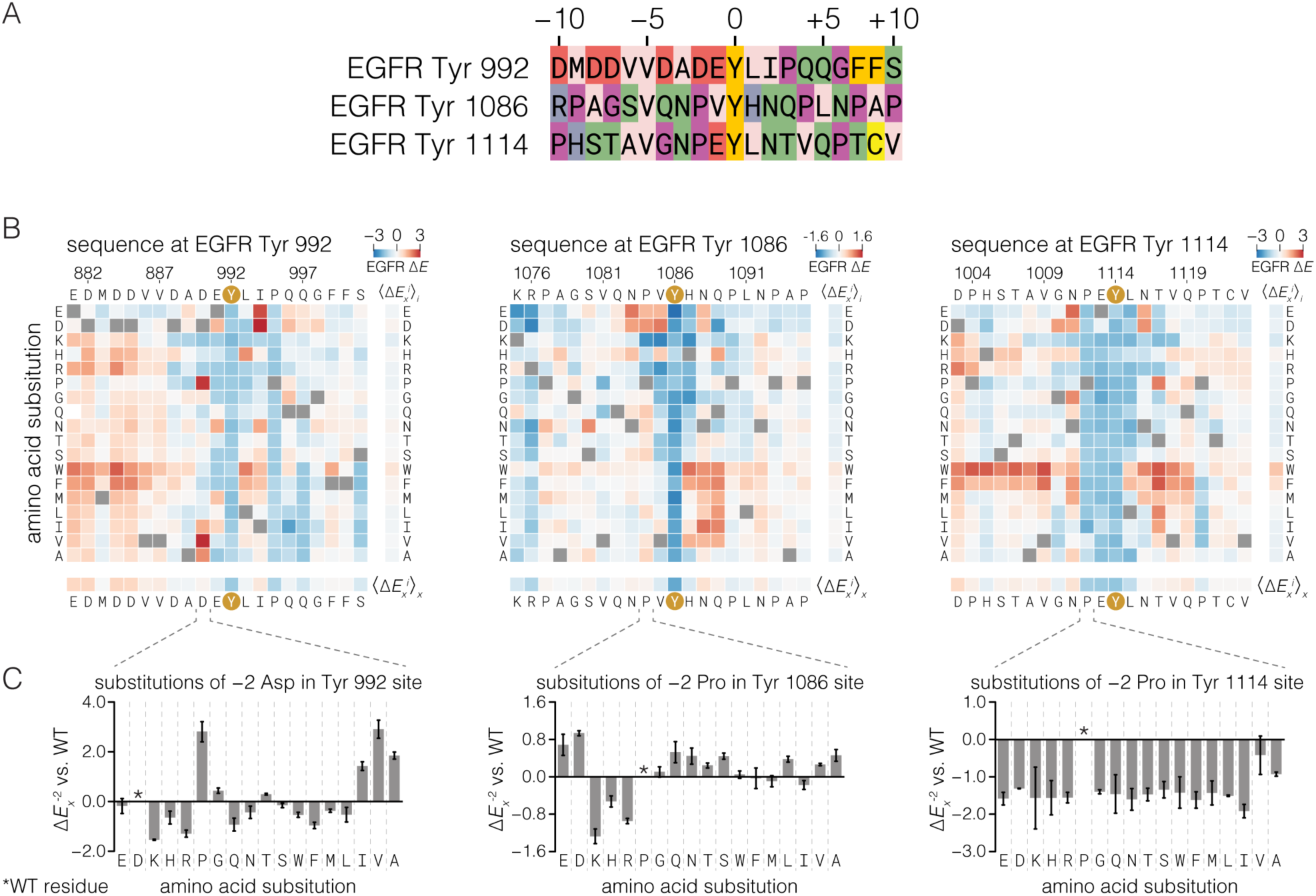
Effect of all single amino acid substitutions on phosphorylation of three EGFR phosphosite peptides by EGFR. A. Sequences of three human EGFR C-terminal tail phosphosites colored by amino acid residue sidechain property (red: acidic; blue: basic; purple: atypical: backbone flexibility; green: polar; tan: aliphatic; gold: aromatic; yellow: cysteine). **B.** Heat maps showing the effect of all single amino acid substitutions (except tyrosine and cysteine) on the phosphorylation level of three EGFR phosphosite peptides relative to wild-type upon phosphorylation by EGFR, measured by bacterial surface-display and deep sequencing. Squares for each substitution *x* of each wild-type position *i* are colored as log-2 fold-enrichment relative to wild-type (Δ*E*_*x*_^*i*^), calculated from read frequency ratios of sorted and input samples. Wild-type residue squares (Δ*E*_*wt*_^*i*^) = 0 by definition) are indicated by gray squares. The Δ*E* scales for each peptide, displayed in the top right corner of each heat map, are not directly comparable because different optimized cell sorting parameters were used for each peptide. Red and blue colors indicate variants that were phosphorylated more or less, respectively, than the wild-type sequence. Row and column mean Δ*E* values are displayed separately. Data are the variant-wise mean of at least two replicates. **C.** Enrichment values for the −2 column (Δ*E*_*x*_^−2^) for each peptide. Error bars, S.E.M.

We used the surface-display/deep sequencing assay to measure the effect of every amino acid substitution along 21-residue stretches spanning these three EGFR phosphosites. The results are presented as heat maps of enrichment values relative to the wild-type peptide for each position in the wild-type sequence (rows) and each substitution to one of 17 other amino acids (columns, excluding tyrosine and cysteine; see Methods) (**Figure 3B**). Mean values across columns and rows are shown as separate bars to the right of and below the main heat maps. The column mean denotes the average effect of perturbing a specific position, whereas the row mean indicates the impact of introducing a specific residue type into the peptide. As expected, mutating the central tyrosine to other residues produces low enrichment relative to the wild-type sequence. Expression levels for each mutant in the Tyr 1114 matrix, measured by cell sorting and deep sequencing, varied marginally when compared to the differences in enrichment scores attributed to phosphorylation (**Supplemental Figure S4**).

A readily apparent feature of all three substitution matrices is the preponderance of positive enrichment values at many different positions. This indicates that the wild-type residues at these positions are not optimal for EGFR phosphorylation. For instance, almost any substitution of an acidic residue 5–10 positions before Tyr 992 increases phosphorylation relative to the wild-type sequence. For the Tyr 992 peptide, these acidic residues are part of an “electrostatic hook” element that is implicated in suppression of kinase activity in the full-length receptor (39, 48, 49). This suggests that the regulatory function of these residues provides an evolutionary constraint on sequence at this phosphosite, at the expense of phosphorylation efficiency.

The three mutational datasets indicate that the EGFR kinase domain does not have sharply defined specificity. In only a few cases is the majority of substitutions at any one position detrimental, notably at the −2, −1, and +1 positions of the Tyr 1114 phosphosite. Substitutions of the −1 or +1 residues away from glutamate and leucine, respectively, in both the Tyr 992 and Tyr 1114 peptides, negatively affect phosphorylation at these sites, in agreement with the emergence of this motif in the Human-pTyr library screen (**Figure 2C**). In further agreement with the Human-pTyr screen, substitutions of the −1 residue by isoleucine either increase phosphorylation (Tyr 992 and Tyr 1086), or reduce it only slightly (Tyr 1114).

### Phosphorylation by EGFR of peptides containing either acidic or large hydrophobic residues at the −1 position may reflect alternate conformations of the bound substrate

The EGFR kinase domain clearly possesses a degree of selectivity at the −1 position of the substrate, as is evident in all three mutational matrices. It is unexpected, however, that both negatively-charged residues (aspartic acid, glutamic acid) and hydrophobic residues (isoleucine, methionine, valine) are permitted at the −1 position in EGFR substrates. A clue to the origin of the dual specificity of EGFR for residues at the −1 position in substrates comes from noting that the effect of replacing the residue at the −2 position depends on whether the residue at the −1 position is acidic or hydrophobic (**Figure 3C**). When the residue at the −1 position is acidic, as in the Tyr 992 and Tyr 1114 phosphosites, then either proline or a residue with a *β*-branched sidechain, such as isoleucine or valine, is strongly preferred at the −2 position. For example, the wild-type Tyr 992 phosphosite has an aspartate at the −2 position, and substitution of this residue by valine or proline leads to increased phosphorylation in the mutational screen, and to increased kinase catalytic efficiency in an in vitro kinase assay using purified proteins and peptides (**Supplementary Figure S4**). If, however, the residue at the −1 position is hydrophobic, as in the Tyr 1086 phosphosite, then there is hardly any constraint on the nature of the residue at the −2 position (**Figure 3C**, middle panel).

We used all-atom molecular dynamics simulations to analyze the conformational space sampled by a substrate peptide. We generated a 200 ns trajectory for an isolated peptide in water, with the sequence of the Tyr 1114 phosphosite, without the kinase domain. In this simulation, the residues at the +1 to +4 positions with respect to the tyrosine were constrained to be in the *β*-conformation, corresponding to how these residues interact with the activation loops of tyrosine kinases (9, 50). We docked instantaneous structures sampled from the simulations on to the EGFR kinase domain (PDB code, 2GS6), using the residues at the +1 to +4 positions as a reference (5). We then examined conformations of the peptide that showed potential interactions with the surface of the kinase domain but did not collide with it.

This analysis showed that are two principal modes in which the residue at the −1 position is recognized by the kinase domain (**Supplemental Figure S5**). In one mode, the residues at the-1 and −2 positions adopt a *β*-conformation. In the other mode, these residues adopt an *α-* conformation. In the *β*-conformation, the tyrosine sidechain and the sidechains at the-1 and-2 position point in alternating directions with respect to the direction of the peptide backbone (**Figure 4A**). An example of this binding mode is provided by the crystal structure of a substrate complex of the c-Kit kinase domain (PDB code 1PKG, (51)). When the-1 and −2 residues of the substrate adopt the *β*-conformation at the EGFR active site, the sidechain of the residue at the −1 position points away from the main tyrosine residue. It is located within a surface pocket that contains the positively-charged sidechains of Lys 855 and Lys 899. In substrates that are primed by phosphorylation at the +1 position, this pocket recognizes the +1 phosphorylated tyrosine residue (26). The sidechain of the −2 residue points toward a hydrophobic pocket near the active site, consisting of Trp 856 and the aliphatic portion of the sidechain of Lys 855. Thus, the *β*-conformation supports binding of substrates with an acidic residue at the −1 position and a hydrophobic residue at the −2 position.

**Figure 4:**
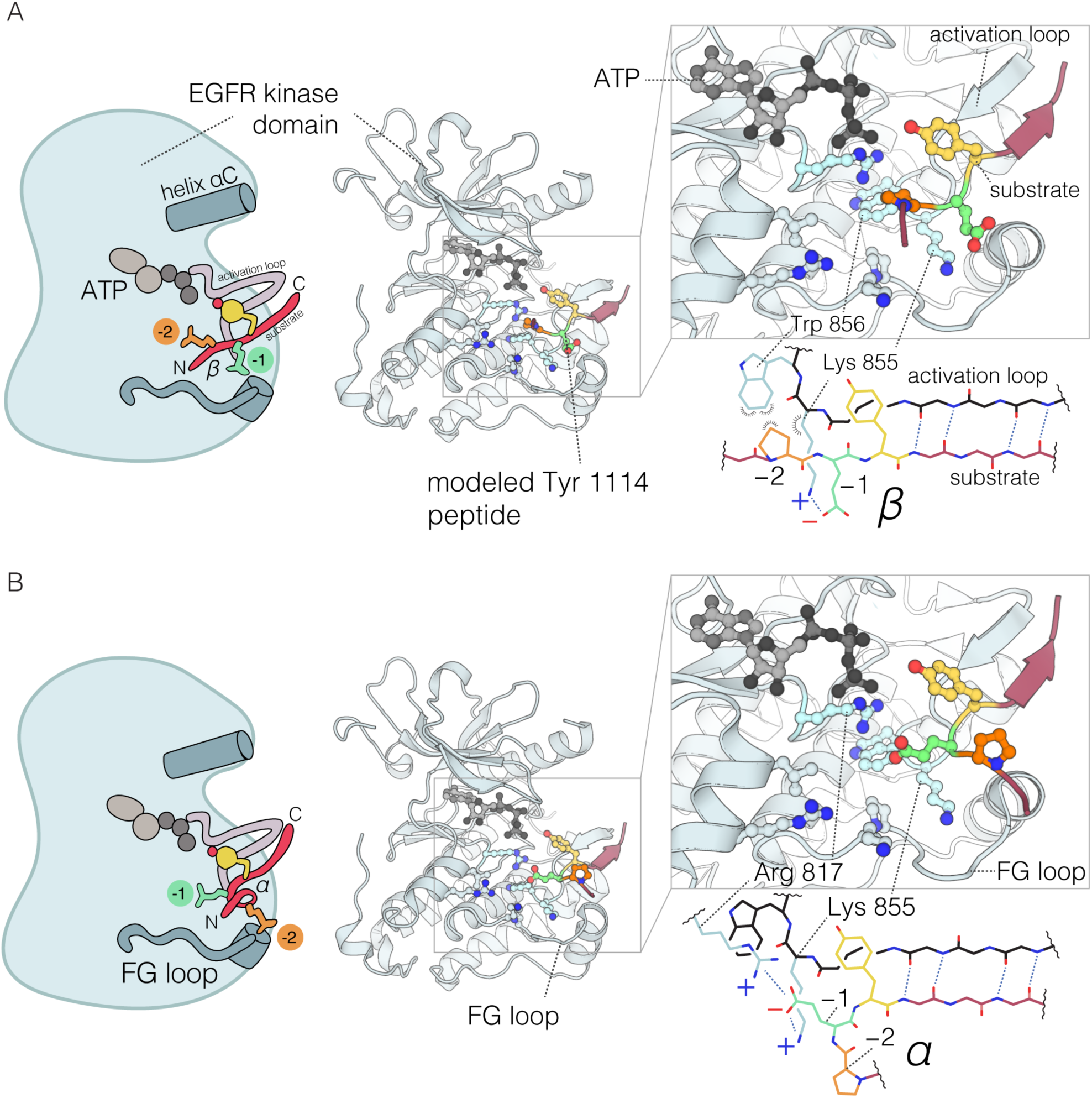
Structural explanation for alternative sequence preferences of EGFR at the −2 position. Selected snapshots from molecular dynamics simulations of an EGFR Tyr 1114 peptide docked onto a peptide-bound crystal structure of the EGFR kinase domain (PDB 2GS6). Two snapshots are shown, with the −1 and −2 residues of the peptide in either the *β* conformation (**A**) or *α* conformation (**B**). Interactions between the −1 and −2 peptide residues and selected residues on the kinase domain are highlighted. Diagrams illustrating the different interactions available between kinase domain residues and a substrate peptide depending on the orientation of the −1 and −2 residues are shown below each zoomed-in view of the active site. A peptide with a −2 Pro and a β conformation of the −1 residue, is diagramed in A, while a peptide with a −2 glutamic acid and an α conformation of the −1 residue is diagramed in B.

Alternatively, when the −1 and −2 residues adopt an *α-*conformation, the sidechain of the residue at the −1 position is pointed in roughly the same direction as the tyrosine residue, towards the same hydrophobic pocket occupied by the −2 residue sidechain in the *β*-conformation. Examples of this binding mode are provided by structures of substrate complexes of EGFR (PDB code 5CZH, (26)) and insulin receptor (PDB code 1IR3 and 1GAG, (50, 52)). The sidechain of the residue at the −2 position is exposed towards solvent (**Figure 4B**). Thus, the *α*-conformation supports recognition by EGFR of peptides with a hydrophobic residue the −1 position, and it does not support sharp discrimination at the −2 position.

The importance of proline at the –2 position in EGFR phosphosites appears to be a consequence of having an acidic, rather than a hydrophobic, residue at the –1 position. We assume that this favors binding of the peptide in a *β*-conformation, which places the residue at the −2 position in the hydrophobic binding site. For the Tyr 1086 phosphosite, which has a hydrophobic residue at the −1 position, the mutational matrix for phosphorylation by c-Src (**Figure 5D**) shows that phosphorylation efficiency is decreased by mutations to either the proline at the −2 position or the valine at the −1 position, suggesting that this peptide is also recognized by c-Src in the *β*-conformation. In contrast to EGFR, c-Src does not tolerate acidic residues the −1 position. A plausible explanation for this effect is the replacement of the positively charged Lys 899 in the FG loop of the EGFR kinase domain by hydrophobic residues in c-Src and other Src-family kinases. The importance of this residue had been noted earlier in an analysis of the differential specificity of the Src-family kinase Lck and ZAP-70 (31).

**Figure 5:**
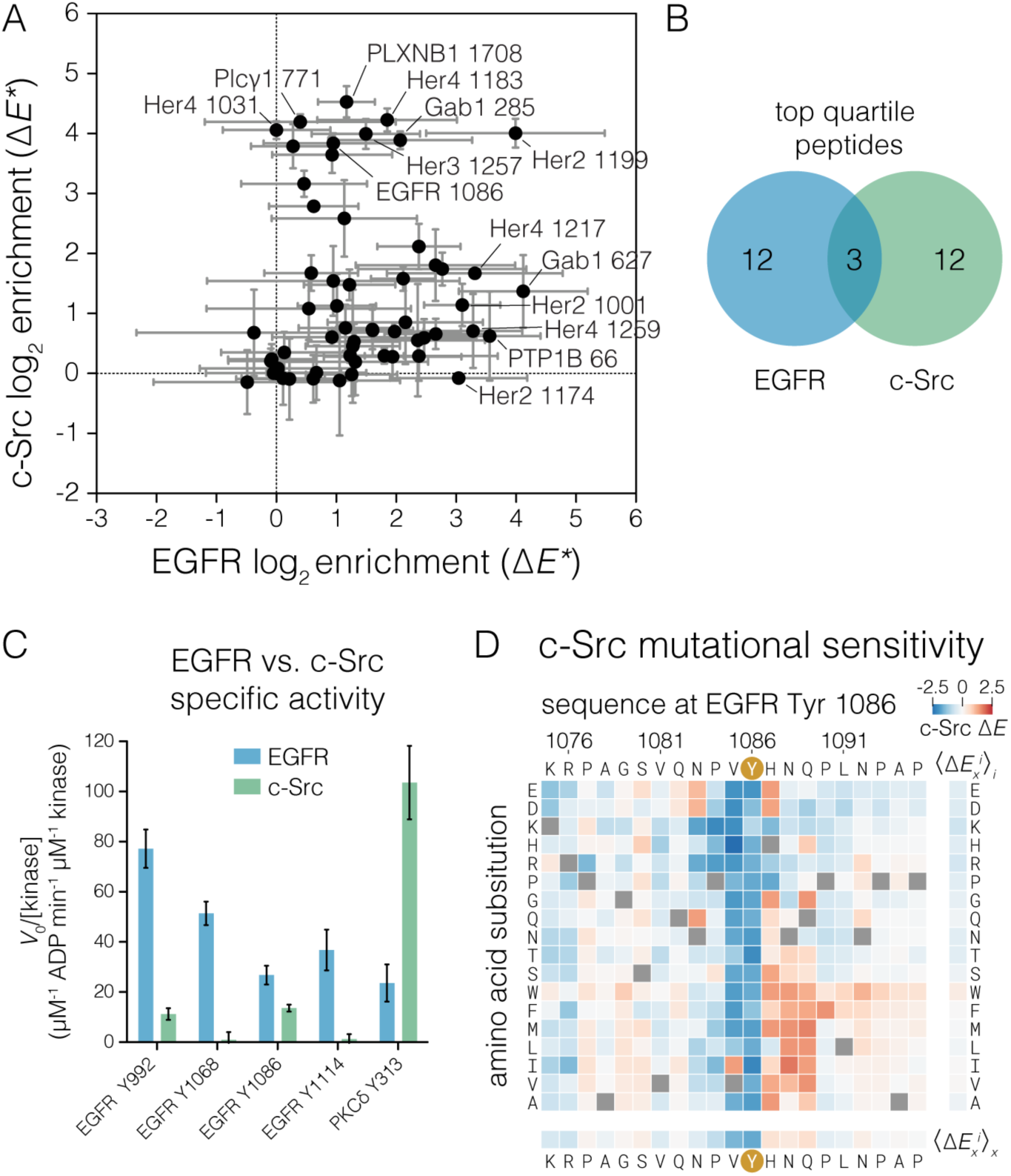
Comparison of c-Src and EGFR specificity with respect to EGFR substrates. A. Relative phosphorylation of EGFR-family phosphosites and reported cytoplasmic EGFR substrates by c-Src versus EGFR. Log-2 fold-enrichment values relative to a non-tyrosine containing control peptide were calculated from peptide read frequencies in sorted and input samples. These enrichment values (denoted *ΔE\**) were corrected by the relative expression level measured for each peptide by cell sorting and deep sequencing. Δ*E*\* values are not comparable on the same scale between kinases. The mean of three replicates with 95% confidence intervals for each kinase is plotted. **B.** Venn diagram showing membership of peptides in the top quartile of Δ*E*\* values for each kinase. **C.** Specific activities measured for c-Src and EGFR by NADH-coupled assay against selected peptides at 0.5 mM peptide. Three EGFR C-terminal tail phosphosites and one c-Src substrate, noted below each set of bars, were measured. Error bars, 95% confidence interval of the mean. **D.** Heat map showing the effect single amino acid substitutions on the phosphorylation level of EGFR Tyr 1086 phosphosite relative to wild-type upon phosphorylation by c-Src, measured by bacterial surface-display and deep sequencing. Δ*E*_*x*_^*i*^ is displayed as heat maps as described in **Figure 2**.

### Tyrosine residues in the tails of EGFR-family members are optimized for selective phosphorylation by EGFR, rather than by c-Src

The fact that EGFR-family phosphosites are highly enriched in sequences that conform to the −1 acidic/+1 hydrophobic rule (**Figure 2D**) suggests that phosphorylation efficiency is indeed an important constraint on the sequences of these sites. If the efficiency of phosphorylation by EGFR is important, then why do the vast majority of these sites exclude a hydrophobic residue at the −1 position, since that would also be consistent with efficient phosphorylation? Src-family kinases and c-Abl, however, efficiently phosphorylate sites with hydrophobic residues at the −1 position (12, 13, 32). Given that c-Src has been shown to participate in EGFR signaling, and therefore is likely to have access to many of the same phosphosites as EGFR-family kinases, we wondered whether minimization of phosphorylation by Src-family kinases and, potentially, other cytoplasmic tyrosine kinases, is a reason for the exclusion of hydrophobic residues at the −1 position in EGFR-family tail phosphosites.

We compared the phosphorylation efficiency of EGFR and c-Src using a focused library of phosphosites in EGFR-family C-terminal tails and reported EGFR substrates (37) using the high-throughput bacterial surface-display assay (**Figure 5A**). For this experiment, we took the additional step of normalizing the enrichment values based on the expression level of each peptide (see **Supplemental Methods**) in order quantitatively compare the two kinases. In this assay, the sites phosphorylated efficiently by EGFR are phosphorylated poorly by c-Src, and vice versa. Except for the Her2 Tyr 1199 peptide, the highly-phosphorylated substrates of EGFR do not include the highly-phosphorylated substrates of c-Src (**Figure 5B**).

To confirm that EGFR phosphosites are poor substrates for Src-family kinases, we compared the catalytic efficiencies of the dimerized EGFR intracellular module and the c-Src kinase domain for selected EGFR phosphosite peptides using an in vitro kinase assay with purified peptides. We compared the specific activity of these kinases at a fixed peptide concentration that is well below the expected values of *K*_M_ for peptide substrates (32, 53, 54). Under such conditions, the catalytic rate is proportional to the catalytic efficiency (*k*_cat_/*K*_M_). For comparison, we also included a peptide from protein kinase Cδ, spanning Tyr-313, which is a good substrate for c-Src (32). Tail phosphosites were phosphorylated by EGFR with efficiency similar to the phosphorylation of a preferred substrate by c-Src, while c-Src phosphorylated the tail sites poorly (**Figure 5C**). This confirms that the c-Src kinase domain is inherently poor at phosphorylating the EGFR tail phosphosites, rather than simply being a less active enzyme than the dimerized EGFR kinase.

The Tyr 1086 phosphosite has valine at the −1 position, which is not optimal for EGFR according to the mutational screen (**Figure 3B**). On the other hand, previous studies using position-specific oriented peptide library screens (13) and bacterial surface-display (32) indicate that c-Src efficiently phosphorylates peptides with a −1 valine. Consistent with this, our results show that Tyr 1086 is among the most efficiently phosphorylated substrates for c-Src in the EGFR tail, but is a relatively poor substrate for EGFR. This observation is also consistent with previous studies reporting that Tyr 1086 is directly phosphorylated by c-Src (27). Intriguingly, mutation of Tyr 1086 in EGFR attenuates the phosphorylation of Tyr 845 in the activation loop of EGFR in a cellular assay, although evidence that this effect is connected to phosphorylation of the EGFR activation loop by c-Src is lacking (49).

We determined the site-specific amino acid substitution sensitivity of the Tyr 1086 phosphosite in EGFR for phosphorylation by c-Src (**Figure 5D**) and compared it to the sensitivity of this phosphosite for phosphorylation by EGFR (**Figure 3B**). Although the Tyr 1086 phosphosite sequence is permissive for phosphorylation by c-Src relative to other EGFR-family phosphosites, this phosphosite is not optimal. Substitution of the histidine residue at the +1 position to most other residues increases phosphorylation by c-Src. A histidine residue at the +1 position of the Tyr 1086 site is a conserved feature of mammal EGFR sequences (**Supplemental Figure S6**), despite having no obvious benefit for EGFR phosphorylation (**Figures 2C, 3A**) or effector protein binding (see below). This suggests that although Tyr 1086 provides a potential channel for phosphorylation of EGFR by c-Src, the sequence of this phosphosite limits the efficiency of such phosphorylation.

### The binding of Shc1 and Grb2 to EGFR phosphosites can be improved by sequence changes that would also increase phosphorylation by c-Src

We expanded the bacterial surface-display method to test the positional amino acid preferences of the SH2 domain of Grb2 and the PTB domain of Shc1, at two EGFR phosphosites, Tyr 1086 and Tyr 1114. We surmised, based on the presence of known binding motifs, as well as binding data from previous studies (19, 46, 55), that the SH2 domain of Grb2 and the PTB domain of Shc1 are prominent adapter proteins that bind to these sites in cells.

In this assay, a mixture of c-Src, c-Abl, and EGFR kinases was used to phosphorylate *E. coli* cells expressing surface-displayed peptide libraries. The phosphorylation was allowed to proceed to completion, as monitored by flow cytometry with an anti-phosphotyrosine antibody (**Supplemental Figure S7**). To assay binding, tandem copies of phosphotyrosine-binding protein domains fused to GFP were used in place of the anti-phosphotyrosine antibody in the flow cytometry selection. Tandem versions of these binding domains were required to maintain a stable signal for the duration of the cell sorting protocol.

The mutational matrices for the binding of the Shc1 PTB domain and the Grb2 SH2 domain (**Figure 6A and 6B**, respectively) have features consistent with previously determined binding motifs for these domains (46). The main determinant of Grb2 SH2 binding is the presence of an asparagine at the +2 position (45, 56). In the mutational matrices, substitution of this residue in both the Tyr 1086 and Tyr 1114 phosphosites is, on average, just as detrimental as substitution of the central tyrosine residue. Similarly, substitution of the asparagine at the −3 position has a uniformly strong negative effect on the binding of the Shc1 PTB at these sites. This corresponds to the known specificity of the Shc1 PTB domain (47, 57). For the Tyr 1086 phosphosite, the −2 proline suggested to be important for PTB binding in earlier studies appears to be dispensable for Shc1 binding, while at the Tyr 1114 phosphosite, the −2 proline appears to be more important.

**Figure 6:**
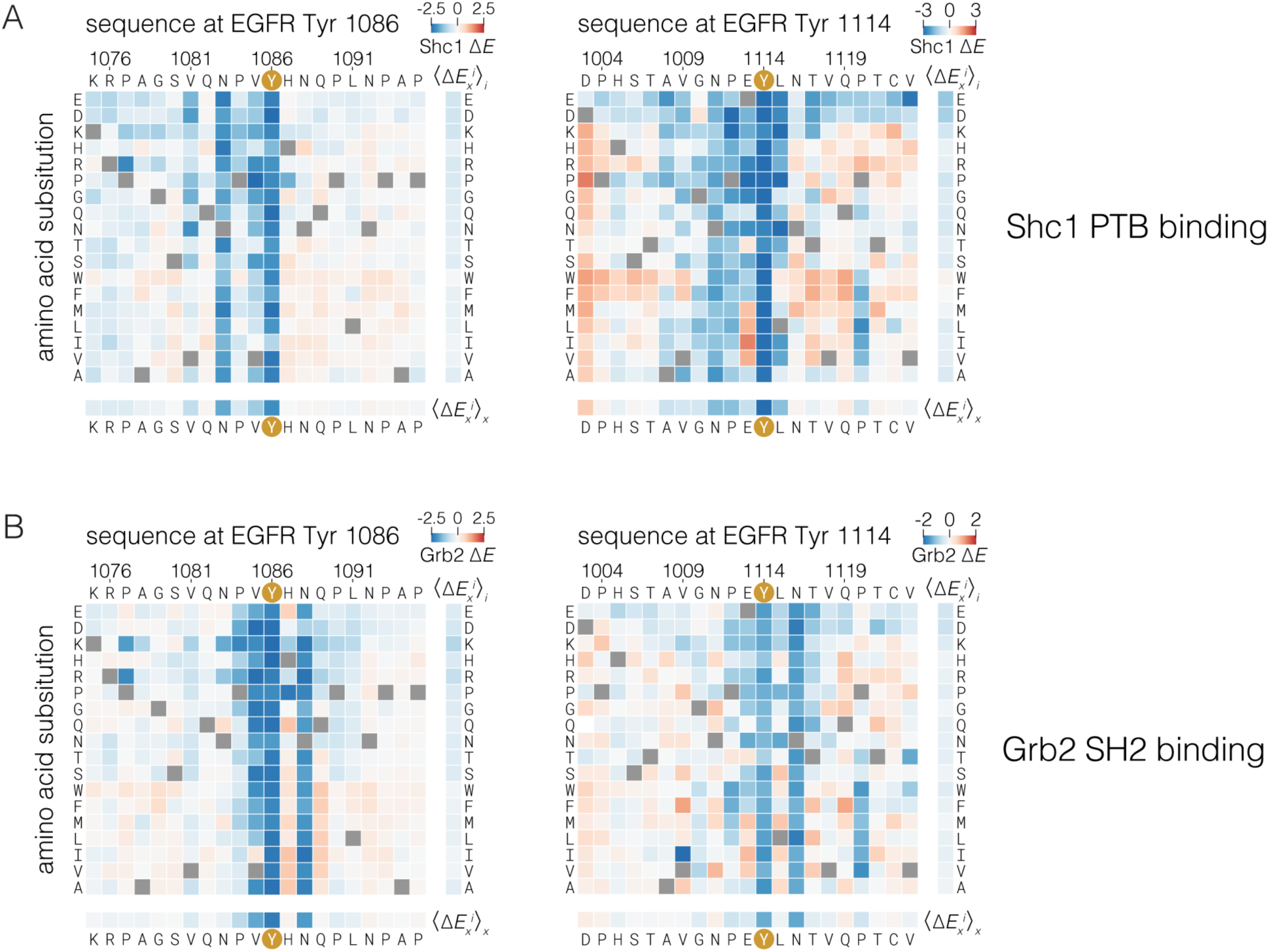
Effect of single amino acid substitutions on the binding of the Grb2 SH2 domain and Shc1 PTB domain to two EGFR phosphosites. Log-2 fold-changes in read frequency ratios relative to wild-type (Δ*E*_*x*_^*i*^) were determined by cell sorting after labeling phosphorylated bacteria displaying peptides with tandem copies of the Shc1 PTB (**A**) or the Grb2 SH2 (**B**) fused to GFP. Δ*E*_*x*_^*i*^ values for single amino acid substitutions are displayed as heat maps, as described in **Figure 2**.

Surprisingly, despite completely different modes for binding to peptides, the Grb2 SH2 and Shc1 PTB domain are both sensitive to substitutions of the −1 valine in the Tyr 1086 phosphosite. Isoleucine, another hydrophobic, *β*-branched residue, is the only substitution that is tolerated by these domains at this position. Introduction of hydrophobic residues, particularly isoleucine or valine, at the −1 position of the Tyr 1114 phosphosite also improves binding, suggesting that a preference for such residues is a shared feature of these domains in multiple sequence contexts. This specificity determinant has not been described previously, and it is not easily explained by available structural models. It is, however, consistent with an alanine-scanning mutagenesis study of phosphosites in the insulin receptor (47).

The preference exhibited by the Grb2 SH2 domain and the Shc1 PTB domain for a hydrophobic residue at the −1 position is at odds with the conserved sequence features found in EGFR-family tail phosphosites, which are not enriched in valine or isoleucine at the −1 position (**Figure 2D**). This preference is, however, aligned with the specificity determinants of Src-family kinases and c-Abl (**Figure 2E and 2F**, (13, 32)). The observation that most EGFR-family tail phosphosites avoid hydrophobic residues at the −1 position, despite such residues being preferred by two of the principal effector proteins that bind to these sites, lends further support to the idea that these sites have evolved to discriminate against phosphorylation by c-Src and, potentially, other cytoplasmic tyrosine kinases.

## Conclusions

The ancestral metazoan organism appears to have had a nearly full complement of cytoplasmic tyrosine kinases that were related closely to their modern counterparts, but orthologs of modern receptor tyrosine kinases were lacking (58, 59). For example, the choanoflagellates, which are among the closest living relatives to true metazoans, have clearly identifiable orthologs of Src-family kinases and Abl. Choanoflagellates also contain numerous receptor tyrosine kinases, but the extracellular domains of these receptors have no counterparts in modern metazoans. Thus, EGFR-family members emerged as important signaling molecules in the background of pre-existing signaling networks that included cytoplasmic tyrosine kinases, such as the Src-family kinases.

This raises the question of whether the tails of the EGFR-family members, which are phosphorylated upon activation of these receptors, are protected from, or perhaps optimized for, phosphorylation by cytoplasmic tyrosine kinases. Our work indicates that evolution has insulated the EGFR tail from efficient phosphorylation by kinases that favor hydrophobic residues at the −1 position, which includes Src-family kinases and c-Abl. While substrates with a hydrophobic residue at −1 can be phosphorylated by EGFR, c-Src, and c-Abl, phosphosites in tails of EGFR-family members generally have an acidic residue at this position. This permits efficient phosphorylation by EGFR, but not by c-Src, and is likely to be an evolutionary adaptation that used negative selection to preserve the integrity of the signals emanating from EGFR family members.

The phosphosites in the tails of EGFR-family members are presented *in cis* for phosphorylation by the receptor, potentially allowing these receptors to rely on proximity rather than sequence specificity for efficient phosphorylation of tail phosphosites. Also, the sequences of EGFR-family tail phosphosites do not conform to the optimal motifs identified for efficient EGFR phosphorylation beyond the −1 and +1 residues (12, 26), making it unclear whether phosphorylation by EGFR is an important parameter constraining the evolution of these sequences. By assaying kinase and binding specificity with respect to discrete, determined sequences, we have discovered that the tail phosphosites do conform to the rules governing efficient phosphorylation by EGFR, but that the sequence motifs found in the tails exclude phosphorylation by Src-family kinases and c-Abl, while maintaining binding of SH2 and PTB domains. Our findings underscore the importance of amino acid sequence context in determining which phosphosites are efficiently recognized by a particular kinase or adapter protein. The context-dependent recognition of proline at the −2 position found in many EGFR-family phosphosites simultaneously allows efficient phosphorylation by EGFR and binding by the Shc1 PTB domain. The relative lack of importance of the +2 position in determining EGFR efficiency allows specification of Grb2 binding, independent of EGFR phosphorylation.

The architecture of the EGFR signaling pathway is organized around a central repertoire of enzymes and adapter proteins that is capable of producing a large variety of cellular outcomes depending on the cellular context (1, 14). The timing and extent of phosphorylation and effector recruitment varies, depending on which ligands and heterodimerization partners are present (60–63). One explanation for the ability of different EGFR-family ligands to produce alternative phosphorylation and effector recruitment patterns is the intrinsic sequence specificity of EGFR-family kinases, which can determine which sites become phosphorylated and able to support effector binding at different thresholds of kinase activity. It will be interesting to see whether intrinsic kinase specificity plays a generally important role in modulating phosphorylation levels in cells, where substrates are often presented to kinases within the locally-dense environment of signaling clusters.

## Methods

### Recombinant Proteins

The sequences of proteins and peptides used in this study are presented in Supplementary Tables 1 and 2, respectively. c-Src kinase domain, tandem Grb2 SH2 and Shc1 PTB GFP fusions, and peptides were expressed in E. coli. The FKBP-and FRB-EGFR intracellular module proteins were expressed in baculovirus. These proteins and peptides purified by standard liquid chromatography methods, as detailed in the Supplemental Methods.

### Bacterial Surface-Display and Deep Sequencing

Quantification of phosphorylation of tyrosine-containing peptides displayed on bacteria was performed largely as described (31). Details for the construction of site-saturating mutagenesis libraries can be found in this reference, and for the construction of the Human-pTyr library can be found in (64). See Supplemental Methods for full details. *E. coli* were transformed with libraries encoding tyrosine-containing peptides at the N-terminus of the engineered surface-display scaffold eCpx. The scaffold also contained a Strep tag at the C-terminus. After induction of peptide expression, cells were harvested and subjected to phosphorylation by a purified kinase, at a concentration, temperature, and duration determined empirically to give ∼30% maximal phosphorylation of a particular library, as judged by flow cytometry with anti-phosphotyrosine 4G10 staining. Cells were sorted by fluorescence activated cell sorting based on anti-phosphotyrosine staining into a single bin corresponding to the brightest 25% of events. DNA from these sorted cells and corresponding unsorted cells was amplified and analyzed by Illumina sequencing in paired mode. The read frequencies for each peptide-coding DNA sequence from the input and sorted samples from each phosphorylation reaction were compared to give an enrichment ratio for each peptide. For site-saturating mutagenesis libraries, the read frequency ratios of mutant peptides were normalized to the that of the wild-type peptide to give a log-relative enrichment value (Δ*E*). For the EGFR substrate library data in **Figure 5A**, this log-relative enrichment was calculated relative to a non-tyrosine-containing control peptide. The data in **Figure 5A** were also corrected for expression level based on relative read frequency ratios calculated from bins sorted on the basis of anti-Strep immunostaining.

Analysis of SH2 and PTB domain binding to the saturation mutagenesis libraries was performed similarly to analysis of phosphorylation, but tandem versions of each binding domain fused to GFP used in place of the anti-phosphotyrosine antibody. The cells used for this analysis were phosphorylated by a mixture of EGFR, c-Src, and c-Abl kinases and monitored for maximal phosphorylation by anti-phosphotyrosine flow cytometry (**Supplemental Figure S7.**)

### Kinase-peptide activity assays

The specific activity of purified kinases against purified peptides was monitored with an enzyme coupled assay based on NADH absorbance (65). Steady-state reaction rates for kinases at 0.5 µM, peptides at 0.5 mM, and at ATP at 0.5 mM at room temperature were converted to specific activity on the basis of NADH absorbance change over time.

### Generation of sequence pLogo of EGFR-family phosphosites

A collection of bona-fide metazoan EGFR-family kinases from was extracted from the EggNOG database (orthology group ENOG410XNSR, (66)), aligned, and filtered by sequence identity of the kinase domain to avoid oversampling taxa that have relatively high numbers of species in the sequence database. C-terminal tyrosine sites were identified from this set of sequences and analyzed for positional amino acid enrichment with the pLogo tool (41), with tyrosines from metazoan transmembrane and intracellular proteins serving as the background distribution. This analysis compares the frequency of each amino acid at each site in a foreground set with the frequency of that amino acid in the same position relative to tyrosine in a background set. The height of each letter corresponds to the log-odds ratio of the binomial probability of observing an amino acid at least as many times in the foreground set as the background set, divided by the probability of observing that amino acid that many times or fewer in the background set.

### Molecular dynamics simulations

The EGFR Tyr 1114 phosphosite peptide was simulated with with NAMD (67) using the CHARMM36 force field (68). The peptide was modeled based on the peptide substrate in the crystal structure of the phosphorylated insulin receptor kinase ((50), PDB 1IR3), and constrained to have residues 1114–1118 adopt *β-*conformation backbone dihedral angles throughout the simulation. After minimization and equilibration of the system with explicit solvent at 300K, simulations were run for 200 ns. Trajectories were analyzed first by clustering frames on the basis of the backbone dihedral angles of residues 1112 and 1113. Representative frames from each cluster were selected visually, and representative frames that do not clash when aligned back onto the EGFR kinase domain are described in **Figure 4.**

## Acknowledgements

We thank Hector Nolla and Alma Valeros of the UC Berkeley Cancer Research Laboratory flow cytometry facility for assistance with cell sorting. We thank Bill Russ and Rama Rangathan at the University of Texas Southwestern Medical Center for assistance with Illumina sequencing and helpful discussions. We thank Xiaoxian Cao of the Kuriyan Lab at UC Berkeley for protein expression assistance. This work was supported in part by grants from the NIH to JK (5R01CA096504-12 and P01AI091580). NHS was supported by the Damon Runyon Cancer Research Foundation postdoctoral fellowship.

## Supplemental Information

Supplemental Figures

**Supplemental Figure S1:**
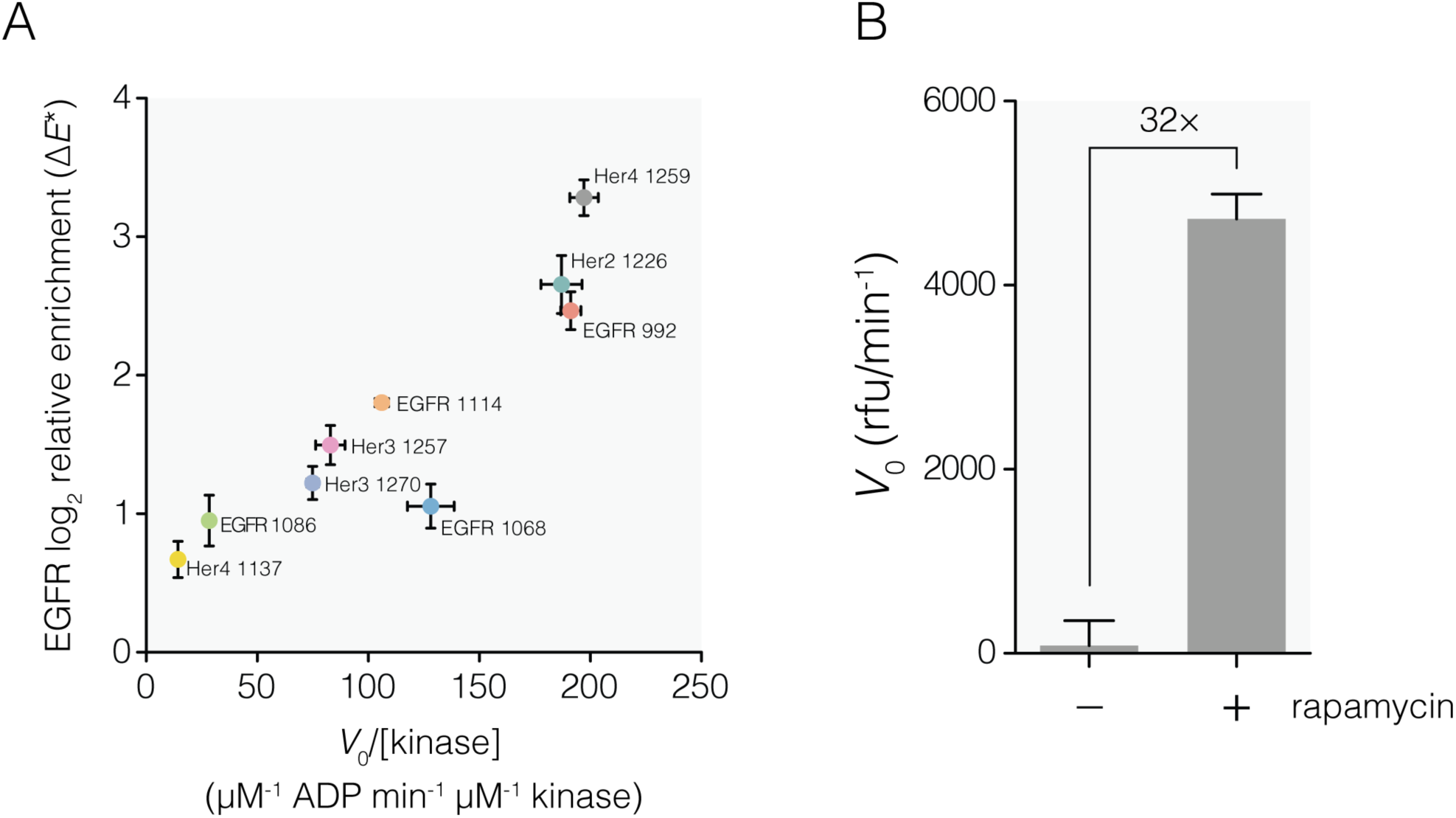
Kinase activity of a dimerized EGFR kinase measured with enzyme-coupled assays and bacterial surface-display coupled with deep sequencing. **A.** The two measurements of enzymatic activity of EGFR for 21-residue peptides corresponding to the indicated EGFR family tail phosphosites were compared. The EGFR protein used in both methods consisted of an equimolar mixture of N-terminal FKBP and FRB fusions of human EGFR residues 663–1186, in the presence of excess rapamycin. The activity on the *x*-axis was measured with a continuous, homogeneous assay wherein the generation of ADP upon phosphorylation of a purified peptide is enzymatically coupled to the oxidation of reduced *β*-nicotinamide adenine dinucleotide (NADH), with a corresponding decrease in absorbance of NADH. Peptides were present at 0.5 mM, below expected *K*_M_ and EGFR dimers were present at 0.2 µM. Activity reported on the y-axis was measured with the bacterial surface-display and deep sequencing assay. For this experiment, peptides were displayed on the surface of E. coli as part of a larger library were subjected to phosphorylation by dimerized EGFR at 0.1 µM dimer for 15 minutes at room temperature, to produce a phosphorylation level of ∼1/3 the maximum obtained by long incubation in the presence of high concentration of kinases. The highly phosphorylated cells were collected by fluorescence activated cell sorting, and the peptide coding portion of the surface-display gene of these cells was sequenced, along with that of the input population. The read frequencies were normalized and plotted as a log-fold-change relative to a negative control peptide containing no Tyr residue and corrected for the separately measured surface-display level. Error bars indicate standard error of the mean from three replicates in each experimental method. B. Effect of forced dimerization on EGFR intracellular module kinase activity. The generic Tyr kinase substrate poly(Glu_4_Tyr)_n_ at 1 mg/ml was subjected to phosphorylation by 50 nM FKBP– and FRB–EGFR dimers in the presence and absence of 1 µM rapamycin. Phosphorylation at various time points was detected with by ADP production enzymatically coupled to production of resorufin. The slope of fluorescence change over time for the linear reaction progress curve is plotted with standard error of the mean, and the fold-increase in rate with the addition of rapamycin is noted.

**Supplemental Figure S2:**
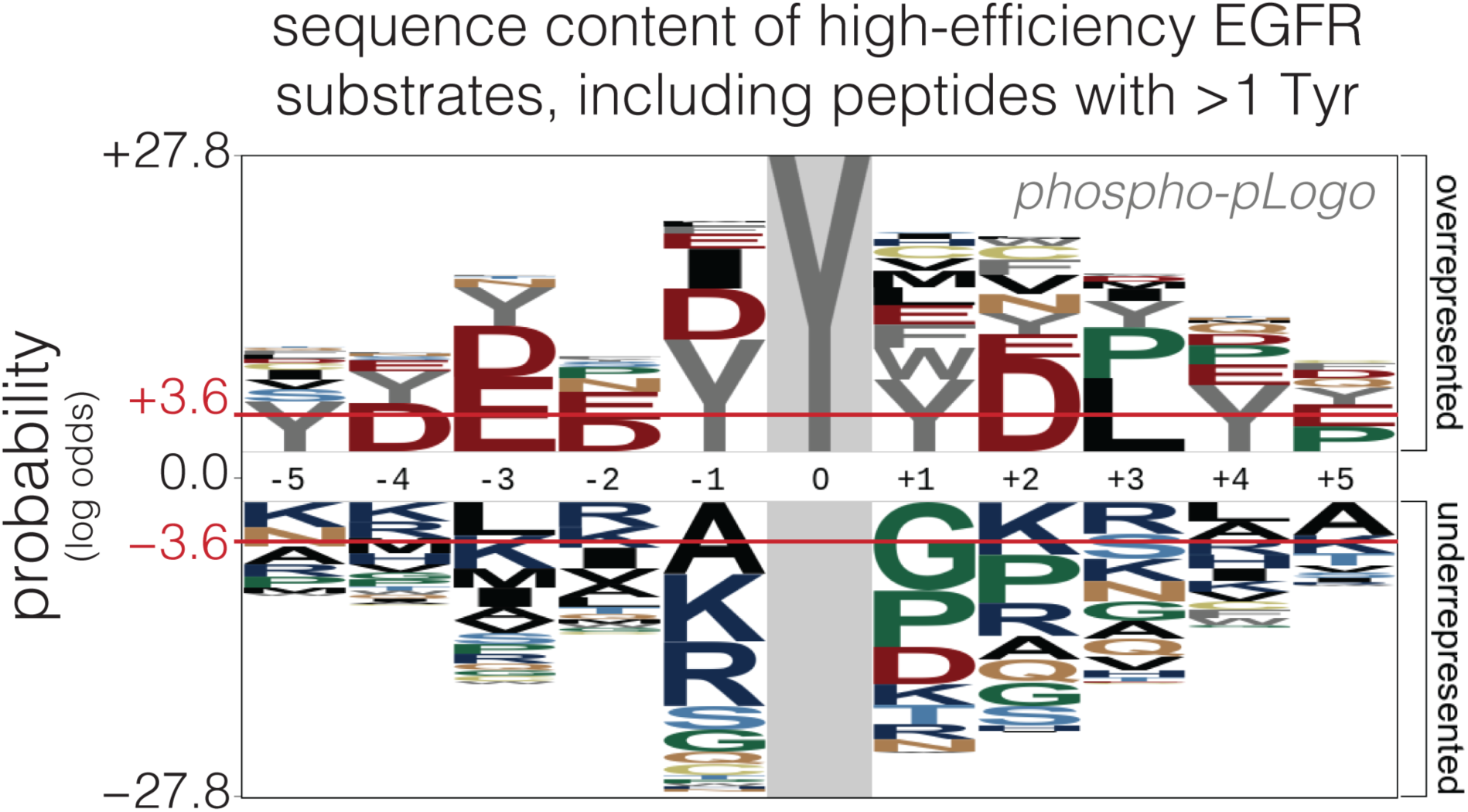
Sequence content of high-efficiency peptide substrates of EGFR from the human proteome, including peptides with more than one Tyr residue. A phospho-pLogo of peptide sequences in the top quartile of read frequency ratios for EGFR, according to the bacterial surface-display/deep sequencing experiment with the Human-pTyr library. This pLogo was generated from the same raw dataset as **Figure 2C**, but including peptides with greater than one Tyr residue in the analysis. Tyr residues appear at multiple positions, but it is not known whether this is a result of multiple Tyr residues becoming phosphorylated and detected by the antibody during the experiment, or due to an improvement of catalytic efficiency for the central Tyr residue when other Tyr residues are present in the peptide.

**Supplemental Figure S3:**
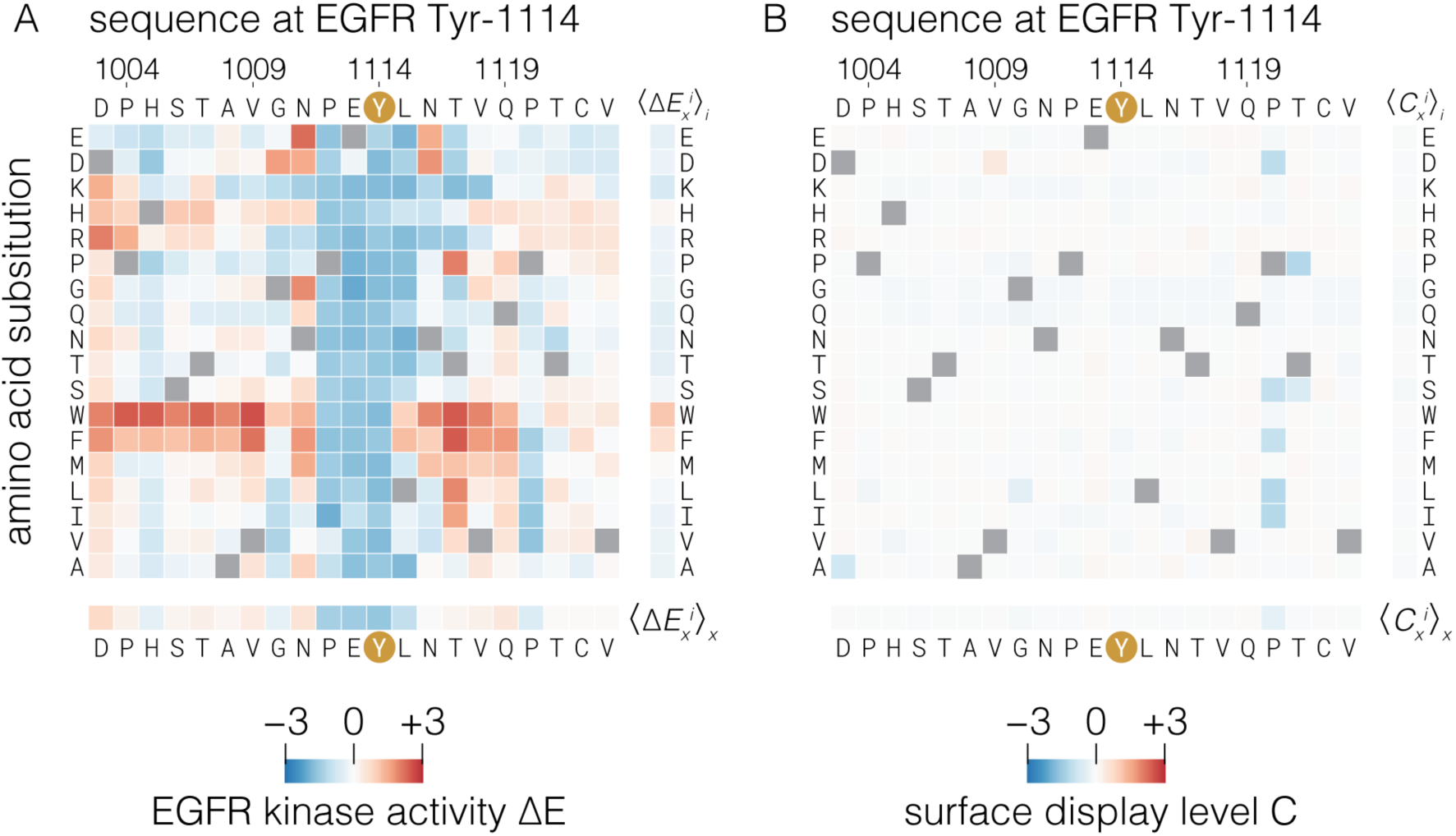
Comparison of relative enrichment differences between variants due to phosphorylation and surface-display level for the Tyr 1114 phosphosite peptide library. The contribution of expression level differences between variants in the calculated enrichment due to phosphorylation by EGFR, Δ*E*, was estimated by measuring the relative surface-display level of each variant in the Tyr 1114 library. Cells displaying the Tyr 1114 library were labeled with a fluorescent anti-Strep tag antibody targeting the surface-display scaffold. These cells were sorted by fluorescence activated cell sorting into six bins spanning the distribution of fluorescence values. The abundance of each peptide in each bin relative to the wild-type peptide was inferred from read frequencies, as measured by Illumina sequencing in the same manner used for phosphorylation level determination. The log-fold differences in expression level relative to the wild-type peptide in the library, *C*_*x*_^*i*^ (**panel B**), were plotted on the same scale as the log-fold differences in phosphorylation level for the Tyr 1114 library (**panel A**, reproduced from **Figure 3B**). White squares indicate minimal differences in expression level for a variant relative to the wild-type peptide, and thus indicate minimal contribution to the phosphorylation enrichment value for that variant, Δ*E*_*x*_^*i*^.

**Supplemental Figure S4:**
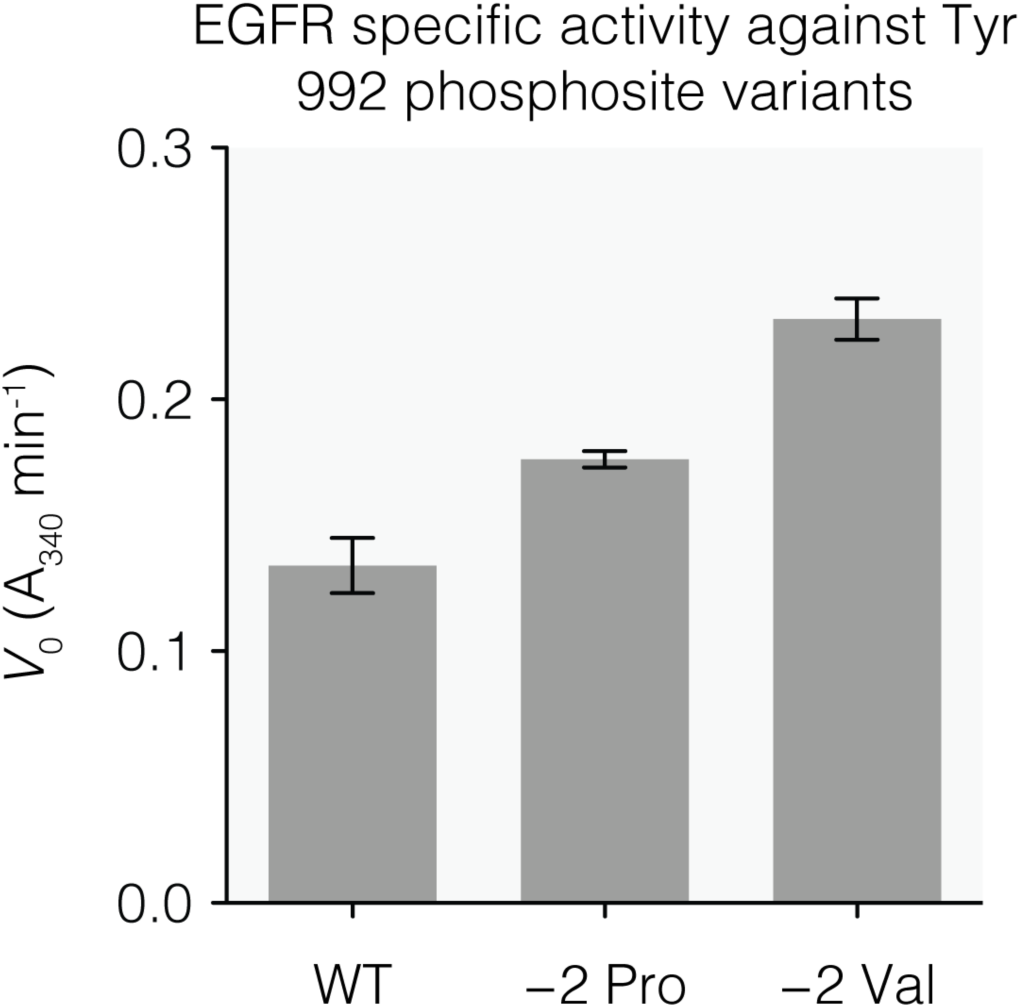
Kinase activity of EGFR against EGFR Tyr 992 phosphosite peptide mutants. Relative kinase activity was measured with an NADH-coupled enzyme assay with purified EGFR intracellular module and purified 21-mer peptides corresponding to the EGFR Tyr 992 phosphosite, with and without the noted substitutions to the wild-type sequence. Steady-state rates were measured in triplicate and plotted as the negative slope of the linear portion of enzyme progress curve of absorbance at 340 nm over time. Error bars, 95% confidence interval.

**Supplemental Figure S5:**
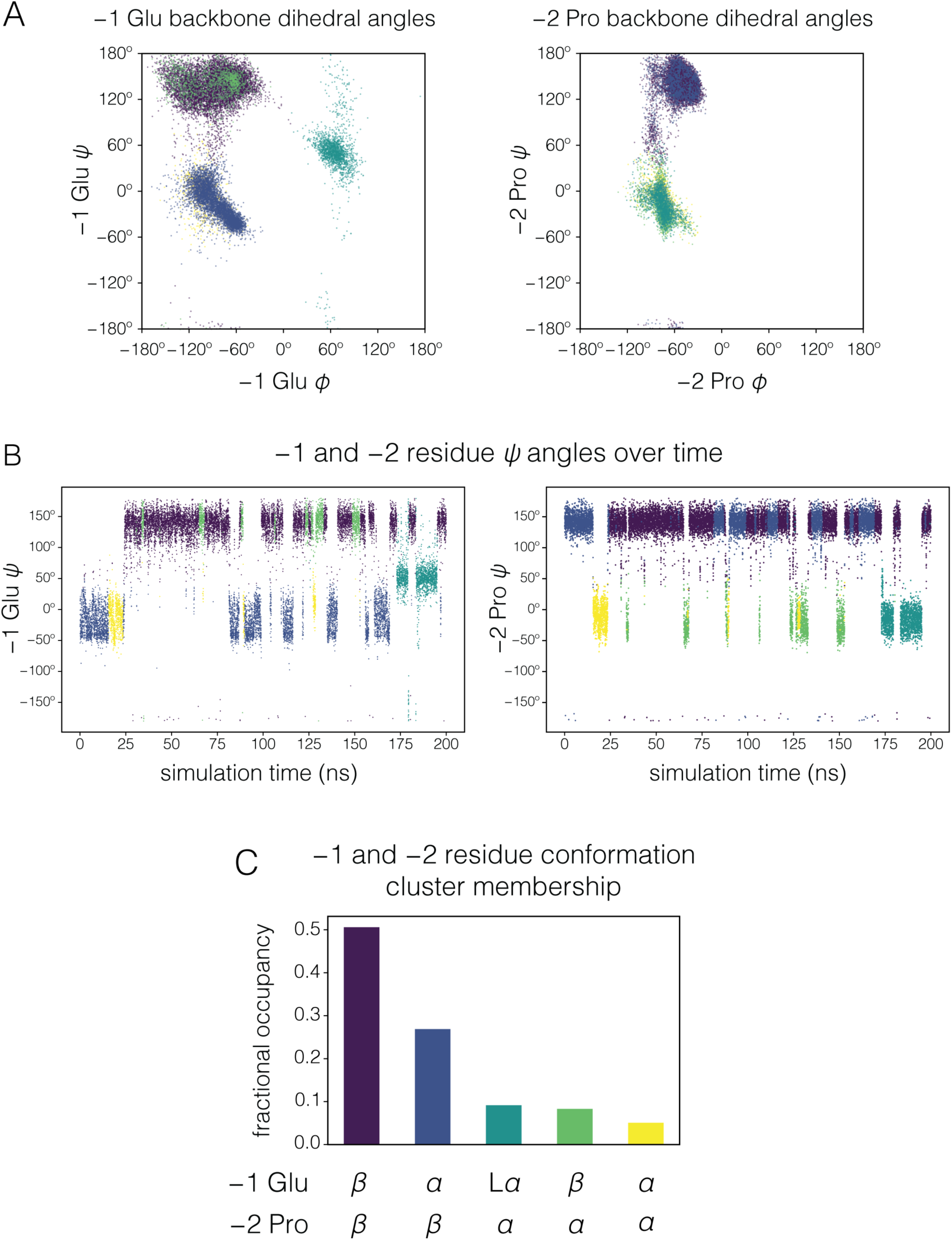
Backbone conformation of the −1 and −2 residues of the EGFR Tyr 1114 phosphosite peptide during molecular dynamics simulations. A. Ramachandran diagrams showing the backbone dihedral angles of the −1 Glu (left panel) and −2 Pro (right panel) residues during a representative molecular dynamics trajectory. Each point represents a frame from the 200 ns trajectory, sampled every 10 picoseconds. The points are colored based on agglomerative clustering performed on four angles, the *φ* and *ψ* angles of the −1 and −2 residues. **B.** *ψ* angles of the −1 and −2 residues over the time course of the simulation, sampled every 10 picoseconds and colored as in A. *ψ* angles between approximately 110° and 180° are considered to represent the *β* conformation, and angles between approximately −50° and 50° are considered to represent the *α* conformation. **C.** Fractional occupancy of the five clusters of −1 and −2 dihedral angles generated for trajectory frames sampled every 10 picoseconds. “L*α*” indicates the left-handed *α*-helical region of the Ramachandran diagram.

**Supplemental Figure S6:**
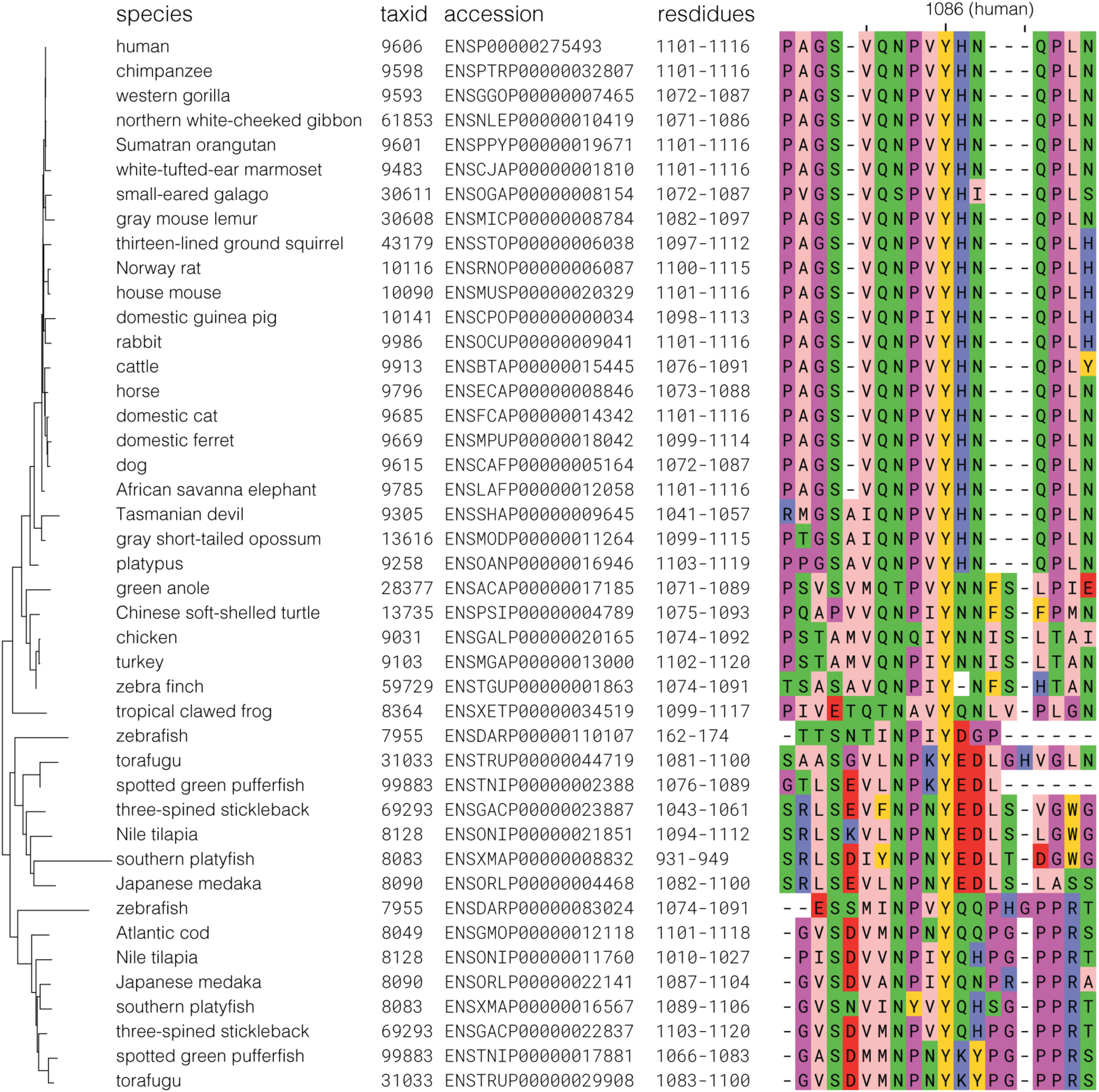
Alignment of EGFR Tyr 1086 phosphosite sequences in the clade after the split between EGFR and Her2. The sequences were aligned with the mafft global homology algorithm and are labeled with common name, NCBI taxid, ENSEMBL translation accession number, and residue boundaries (including signal sequences). The phylogenetic relationship for the corresponding full-length EGFR sequences, taken from the EggNOG database, is shown on the left. A His residue at the +1 position of the Tyr 1086 phosphosite is a conserved feature of mammalian EGFR sequences. No phosphosites contain a −1 acidic or +1 hydrophobic residue.

**Supplemental Figure S7:**
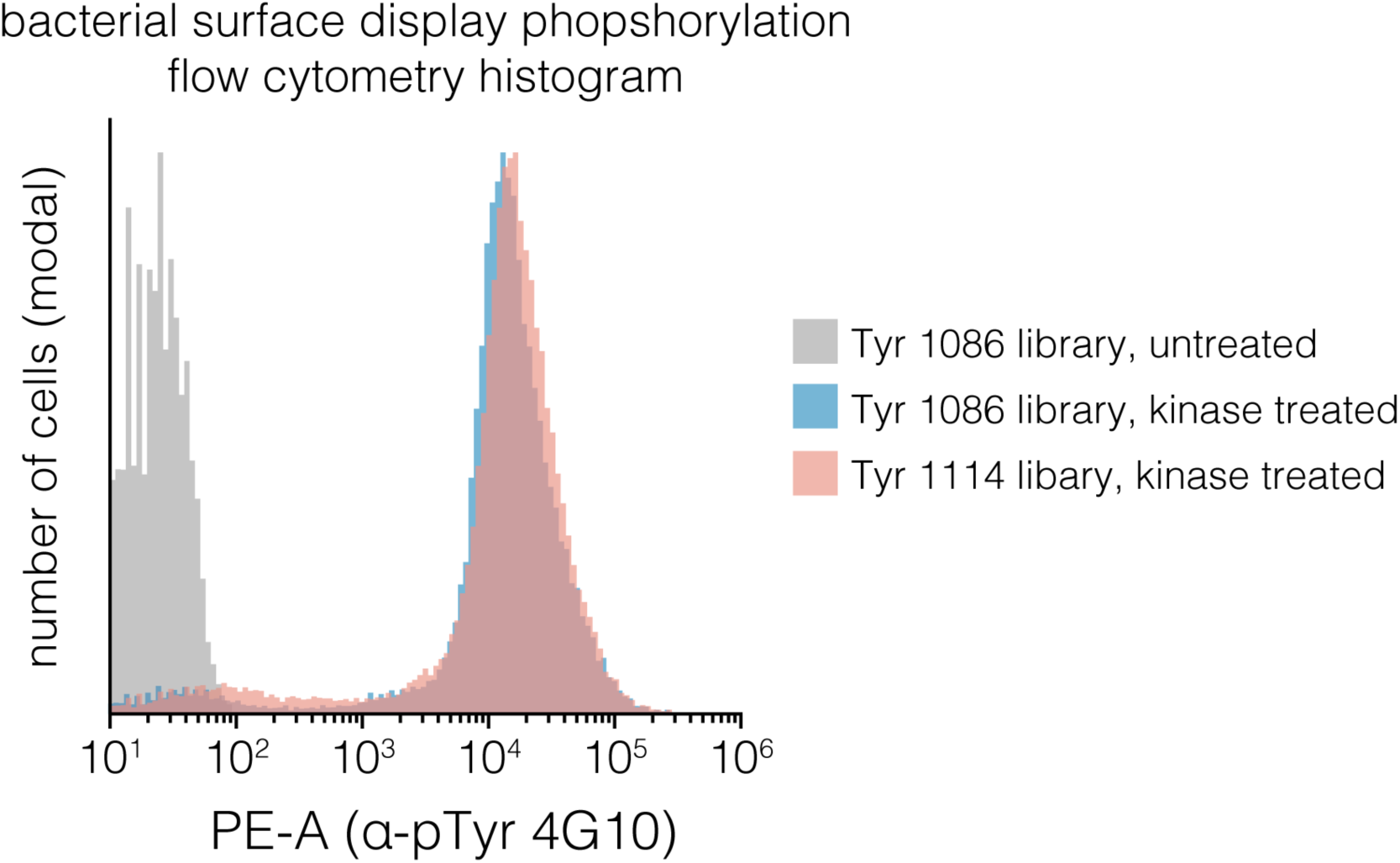
Flow cytometry histogram of fully-phosphorylated bacteria displaying mutagenesis libraries. A sample of the bacteria that served as an input to the surface-display binding experiments presented in **Figure 6** were stained with an anti-phosphotyrosine antibody (4G10) and analyzed by flow cytometry. Site-saturation mutagenesis libraries corrsesponding to the EGFR Tyr 1086 and 1114 phosphosites were either treated with a mixture of EGFR, c-Src, and c-Abl kinases for 1 hour at room temperature (“kinase treated”) or incubated in the absence of kinases (“untreated”). The single, narrow main peaks in the histogram indicate the libraries wer uniformly phosphorylated.

## Supplemental Methods

### Recombinant Proteins

Amino acid sequences for the proteins used in this study are listed in **Supplementary Table S1.**

#### c-Src kinase domain

The chicken c-Src kinase domain (corresponding to residues 257–525 human c-Src) was expressed and purified as previously described (32, 69).

#### FKBP-and FRB-EGFR intracellular module fusions

Constructs consisted of a non-cleavable His_10_ tag followed by either human FKBP1A (residues 3–108) or the FRB domain of human mTOR (residues 2018–2112). These were fused N-terminally to human EGFR residues 663–1186, including the C-terminal part of the juxtamembrane element and the full C-terminal tail. The construct terminated with a FlAsH-tag binding sequence (CCPGCC), which was not utilized in this study. These EGFR constructs were inserted into the pFastBac1 vector and expressed in Sf9 cells with the Bac-to-Bac system (ThermoFisher Scientific), in ESF 921 medium (Expression Systems). The EGFR proteins were purified by TALON Co^2+^ affinity, anion exchange, and size exclusion chromatography. The proteins were concentrated and stored in a buffer containing 50 mM Tris pH 8.0, 150 mM NaCl, 10% glycerol, and 1 mM TCEP.

#### Peptides

EGFR tyrosine-containing peptides were produced recombinantly in *E. coli*. Coding sequences for 21-residue peptides were inserted C-terminal to yeast SMT3 with an N-terminal His_6_ tag in a pET-derived vector. The proteins were purified with Ni^2+^-affinity and anion exchange chromatography. The SUMO moiety was removed with yeast Ulp1 protease, and cleaved peptides were isolated with reverse-phase HPLC in water with 0.1% TFA, with elution by a gradient of acetonitrile with 0.1% TFA. Peptides were lyophilized, resuspended in water, and dialyzed against 100 mM HEPES pH 7.5. Peptide concentrations were determined by Tyr absorbance at 275 nM (1410 M^−1^ cm^−1^ extinction coefficient). The Protein Kinase C Tyr 313 15-mer peptide was purchased from Elim Biopharmaceuticals.

#### Tandem SH2 and PTB domain GFP fusions

The tandem human Grb2 SH2 (residues 55 to 152) and human Shc1 PTB domain (residues 147 to 318) DNA sequences were constructed by overlap extension PCR and inserted into a pET-based vector, with an N-terminal His_6_-tag and TEV protease site and C-terminal eGFP. The two copies of each binding domain were connected by a 20-residue Gly/Ser linker, and these were connected with a 10-residue linker to GFP. These proteins were expressed in *E. coli* and purified using Ni^2+^-affinity, anion exchange, and size exclusion chromatography. His_6_ tags were removed prior to size exclusion chromatography. Protein concentration was determined by absorbance of GFP at 488 nm.

### Bacterial Surface-display and Deep Sequencing

#### Human-pTyr library phosphorylation analysis

The specificity profile for the dimerized EGFR intracellular module against the Human-pTyr library of phosphosites was determined as described by (32). Details of the library construction and contents can be found in this reference. Briefly, *E. coli* (strain MC1061) expressing the library were subjected to phosphorylation by 0.5 µM dimerized EGFR intracellular module for 15 minutes at room temperature. These conditions were determined to produce a median phosphorylation level of ∼30% compared to a fully phosphorylated sample, as judged by flow cytometry with staining with Milli-Mark anti-Phosphotyrosine 4G10 phycoerythrin antibody conjugate (Millipore Sigma). The top 25% brightest cells in the PE channel were sorted on a BD Influx cell sorter. ∼4 million sorted and unsorted cells were harvested by centrifugation and boiled to release DNA. The peptide-coding portion of the surface-display plasmid was PCR amplified in two steps to append Illumina indices and adapters. The samples were multiplexed and sequenced in paired-end mode on an Illumina MiSeq sequencer. Paired-end reads were assembled, trimmed, and mapped to peptide sequences in the library. Peptide sequence read frequency in each sample was calculated as the number of reads for a given peptide divided by the total number of aligned reads for that sample. The ratio of read frequency of each peptide in the sorted sample divided by the frequency of the same peptide in the unsorted sample gives the read frequency ratio plotted and analyzed in **Figure 2**. For the data presented in **Figure 2**, for all kinases, only peptides containing a single Tyr residue were analyzed. Phosphosite sequences with two tyrosines had the non-central Tyr mutated to alanine during library construction. Phospho-pLogo diagrams were generated as follows. The central 11 residues of the phosphosite sequences (five residues before and after Tyr) with read frequency ratios in the top 25% of ratios within a given sample across at least two replicate samples (top 25% in replicate one AND replicate two) were counted as highly phosphorylated sequences. These highly phosphorylated sequences were used as the foreground set in the calculation of a pLogo, as described in (41). The background for this calculation set was the set of single-Tyr sequences observed in the unsorted sample. This analysis was repeated using the raw read frequency ratio data for chicken c-Src and human c-Abl published in (32).

#### Construction of phosphosite single-site saturation mutagenesis libraries

Single-site saturating mutagenesis libraries were constructed as described in (31). Briefly, oligonucleotides containing one degenerate NNS codon each (Integrated DNA Technologies) were used in an overlap-extension PCR reaction to generate constructs with a 21-residue peptide-coding sequence in-frame with the eCpx bacterial surface-display scaffold (70), with one mutated sequence position per DNA fragment. These fragments were pooled, restriction digested, and ligated into the SfiI sites of the pBAD33 vector. The pooled, ligated mixture was transformed into TOP10 *E. coli* (ThermoFisher Scientific) by electroporation, and library DNA was isolated with a silica membrane spin column (Zymo Research).

#### Phosphorylation analysis of phosphosite single-site saturation mutagenesis libraries

Libraries were phosphorylated as described for the Human-pTyr library, at room temperature, with kinase concentrations and durations that produced ∼30% of maximal library phosphorylation as judged by flow cyotmetry. Cells were labeled with anti-phosphotyrosine 4G10 antibody-PE conjugate, and the top 15% of cells in the PE channel were sorted on a BD FACSAria Fusion cell sorter. Sorted and input cell samples were sequenced as described for the Human-pTyr library, above. Relative enrichment for each sequence position *i* mutated to each amino acid substitution *x* versus the wild-type (WT) variant, Δ*E*_*x*_^*i*^, was calculated from the read frequencies in the sorted and unsorted samples with the following formula:

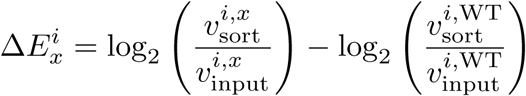

where *v*_sample_^*i,x*^ is a read frequency of variant *x* at position *i* for a particular sample. Data in **Figures 3 and 5** are the mean from at least two biological replicates.

#### Phosphorylation analysis of EGFR substrate phosphosite library, with expression level correction

A collection of 21-residue single-tyrosine peptides corresponding to human EGFR-family C-terminal Tyr residues and EGFR substrates reported in the PhosphositePlus Database (37) was assembled by overlap-extension PCR and inserted into the eCpx scaffold. The sequences are listed in **Supplementary Table S2**. This library was expressed, phosphorylated by the dimerized EGFR intracellular module or c-Src kinase domain, sorted, and sequenced as described for the Human-pTyr library. Enrichment ratios for a peptide *p* relative to a peptide containing no Tyr residues, Δ*E*_*p*_, were determined for each peptide with the following formula:

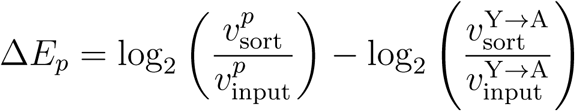

where *v*_sample_^Y →A^ denotes the read frequency of a peptide corresponding the EGFR Tyr 1173 phosphosite, with the tyrosine mutated to alanine, in a particular sample. Relative expression levels were also measured for each peptide in order to correct for expression level differences that could show up as phosphorylation level differences in the bacterial surface-display/deep sequencing assay. Expression levels were monitored in a separate experiment using a Strep-tag at the C-terminus of the eCpx scaffold, detected with the StrepMAB-Classic chromeo 488 antibody conjugate (IBA Lifesciences). Cells were sorted into 6 bins based on fluorescence in the FITC channel with a BD FACSAria Fusion cell sorter, and read frequencies were calculated for each peptide *p* in each bin. These read frequencies were weighted by the number of flow cytometry events *e* in each bin versus the total number of events, as in

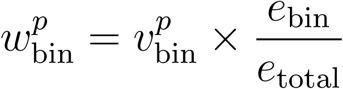

The resulting weighted frequencies for each peptide in each bin, *w*_bin_^*p*^, versus the log-transformed mean fluorescence measured for all flow cytometry events in each bin were fitted to a simple Gaussian distribution, giving an estimated weighted mean fluorescence ⟨*w*^*p*^ ⟩ for each peptide. This value was compared to that for the no-tyrosine control peptide and the measured mean fluorescence of the lowest bin, *w*_0_^*p*^ to generate a relative expression level *Cp*, according to the following formula:

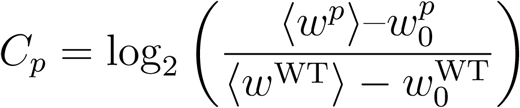

Finally, the expression-corrected relative enrichment ratio for each peptide, Δ*E*_*p*_*, was calculated as:

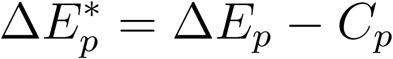

Δ*E*\* values for each peptide in the library, calculated for EGFR and c-Src, are plotted in **Figure 4C.** Relative expression level for each mutant versus the wild-type peptide were measured for the Tyr 1114 phosphosite mutagenesis library by the same method, with the wild-type peptide serving as the reference, and plotted in **Supplemental Figure S3.**

#### Binding analysis of phosphosite single-site saturation mutagenesis libraries

Libraries were phosphorylated as described for the Human-pTyr library, but with a mixture of 2.5 µM c-Abl kinase domain, 2.5 µM c-Src kinase domain, and 1 µM dimerized EGFR intracellular module (including 2 µM rapamycin). These preparative phosphorylation reactions were incubated for 1 hour at room temperature. They also included the addition of 50 µg/ml rabbit muscle creatine phosphokinase and 5 mM creatine phosphate (Sigma Aldrich) to regenerate ATP. To confirm complete library phosphorylation, a sample of cells treated in this manner was monitored by flow cytometry based on anti-phosphotyrosine-PE labeling. The phosphorylated cells were harvested by gentle centrifugation and washed once with binding buffer containing 50 mM HEPES pH 7.5, 150 mM NaCl, 1 mM TCEP, and 0.2% BSA. Cells were then resuspended in this buffer containing either 5 µM tandem Grb2 SH2-GFP or 1 µM tandem Shc1 PTB-GFP and incubated for 1 hour at room temperature. After labeling, the cells were centrifuged, and the label-containing supernatant was discarded. The cells were washed in this manner one time with labeling buffer and finally resuspended for fluorescence-activated cell sorting. Cells were sorted by GFP fluorescence on a BD FACSAria Fusion, with the top 15% of cells in the FITC channel collected. DNA from the sorted and input cell samples was amplified, indexed, and sequenced as described for the phosphorylation experiments, above. Position-wise relative enrichment for each mutant versus wild-type, Δ*E*_*x*_^*i*^, was calculated in the same manner as for mutagenesis matrices with respect to phosphorylation.

### Kinase-peptide Activity Assays

Kinase activity was measured with an enzyme coupled assay based on NADH absorbance (65). Kinases were assayed at 0.5 µM enzyme concentration in a buffer containing 50 mM HEPES pH 7.5, 50 mM NaCl, 10 mM MgCl_2_, 1 mM TCEP, 1 mM phosphoenol pyruvate, 0.5 mg/ml NADH,∼40 U/ml pyruvate kinase, and ∼60 U/ml lactate dehydrogenase (from rabbit muscle, Sigma Aldrich). Reactions contained either 0.5 mM or no peptide. Reactions containing EGFR also contained 1 µM rapamycin (LC Laboratories). Reactions were initiated with the addition of ATP to a final concentration of 0.5 mM. Reaction progress was monitored by the change in absorbance at 340 nm over time, at 25° C. Kinase activity was calculated as the difference in rates between reactions with and without substrate peptide, using the extinction coefficient of NADH at 340 nm of 6220 M^−1^ cm^−1^.

### Bioinformatics

#### Generation of sequence pLogo of EGFR-family phosphosites

EGFR-family C-terminal tail phosphosite sequences were collected from the EggNOG database (orthology group ENOG410XNSR, (66)) and filtered to include only the sequences in the Eumetazoa and Porifera taxonomic groups (NCBI taxids 6072 and 6040). Full-length protein sequences were further filtered to exclude sequences with greater than 90% sequence identity within the kinase domain, in order to obtain a set of sequences that evenly samples natural evolutionary sequence space, without oversampling taxa that have relatively high numbers of species in the sequence database. Kinase domain boundaries were identified by querying each EGFR sequence against the SMART database (72). These domain sequences were aligned with Mafft (G-INS-I algorithm, (73)). and then filtered by sequence identity with the CD-HIT web service (74). From the resulting set of full-length sequences, phosphosite sequences were extracted as the five residues before and after each Tyr that occurs after each kinase domain, as identified above. This list of 11-residue sequences was used as the foreground set in the pLogo web service (41) to generate the sequence pLogo shown in **Figure 2D**. The background set was a random sampling (to keep the total number of sequences below the web server’s limit) of the UniRef50 database of representative sequences (38), filtered to include verified metazoan proteins, tagged as having either transmembrane or intracellular localization.

#### Sequence alignment of EGFR-family phosphosites

Full-length sequences in the EGFR branch of the metazoan EGFR-family (66) were roughly aligned with Mafft (73), and the Tyr 1086 site was identified visually in each sequence with Jalview (71). These sites were then realigned with Mafft and ordered based on the topology of the EGFR-family tree in the EggNOG database (66).

### Molecular Dynamics Simulations

Simulations were prepared with VMD (75) and generated with NAMD (67) using the CHARMM36 force field (68). For all stages, electrostatics were calculated with the particle mesh Ewald summation, and a non-bonded cutoff of 12 Å was employed. A starting model was generated for the peptide corresponding to human EGFR residues 1110–1118 by assigning backbone and C*β* atoms of residues 1114–1118 to the coordinates of chain B of the crystal structure of the insulin receptor kinase domain bound to a substrate peptide ((50), PDB 1IR3). Residues 1110–1113 were modeled in an extended conformation. The peptide model’s N-and C-termini were capped with N-acetyl and N-methylcarboxamide groups, respectively. This model was solvated in a rectangular box with TIP3P water, and with sodium and chloride ions for an effective ionic strength of 150 mM.

The model was minimized and equilibrated as follows. 1000 steps of conjugate gradient minimization were performed with the protein residues fixed and solvent molecules allowed to move. This was followed by 1000 steps of conjugate gradient minimization in which the peptide atoms were allowed to move, but with an additional sinusoidal potential with a spring constant of 10 kcal mol^−1^ radian^−2^ applied to the backbone *φ* angles of residues 1115–1118 and *ψ* angles of residues 1114–1117, to keep these residues in their starting *β-*strand conformation. After minimization, the system was assigned random velocities and equilibrated under constant number, temperature, and pressure (NPT) conditions at 300 K for 50000 steps, at 2 fs step^−1^ (for 1 ns total), with all protein atoms restrained by a harmonic potential, with a spring constant of 1 kcal mol^−1^ Å^−2^. This was proceeded by production simulation under the same conditions, with all atoms unrestrained except for the dihedral restraints for residues 1114–1118 described above. (The first 1 ns was discarded from analyses to allow for additional equilibration.)

Simulations were analyzed with the cpptraj package of the Amber software suite (76) and pymol (77). Simulation frames were aligned to the EGFR kinase domain as follows. First, the co-crystal structure of the EGFR kinase domain and a bi-substrate analog ((5), PDB 2GS6) was aligned based on kinase domain backbone atoms to the insulin receptor kinase domain bound to substrate ((50), PDB 1IR3). Then, simulation frames were aligned to the substrate peptide of the insulin receptor crystal structure based on the backbone atoms of residues 1114–1117 of the simulated EGFR peptide, producing models of the EGFR kinase domain bound to a peptide substrate. Representative snapshots for **Figure 3** were chosen by visual inspection of sets of simulation frames obtained by clustering based on the backbone dihedral angles of residues 1112 and 1113. Frames, sampled every 10 ps, were assigned to one of five clusters based on the backbone dihedral angles of the –1 and –2 residues of each frame, by agglomerative clustering in a custom python script.

**Supplemental Table S1:**
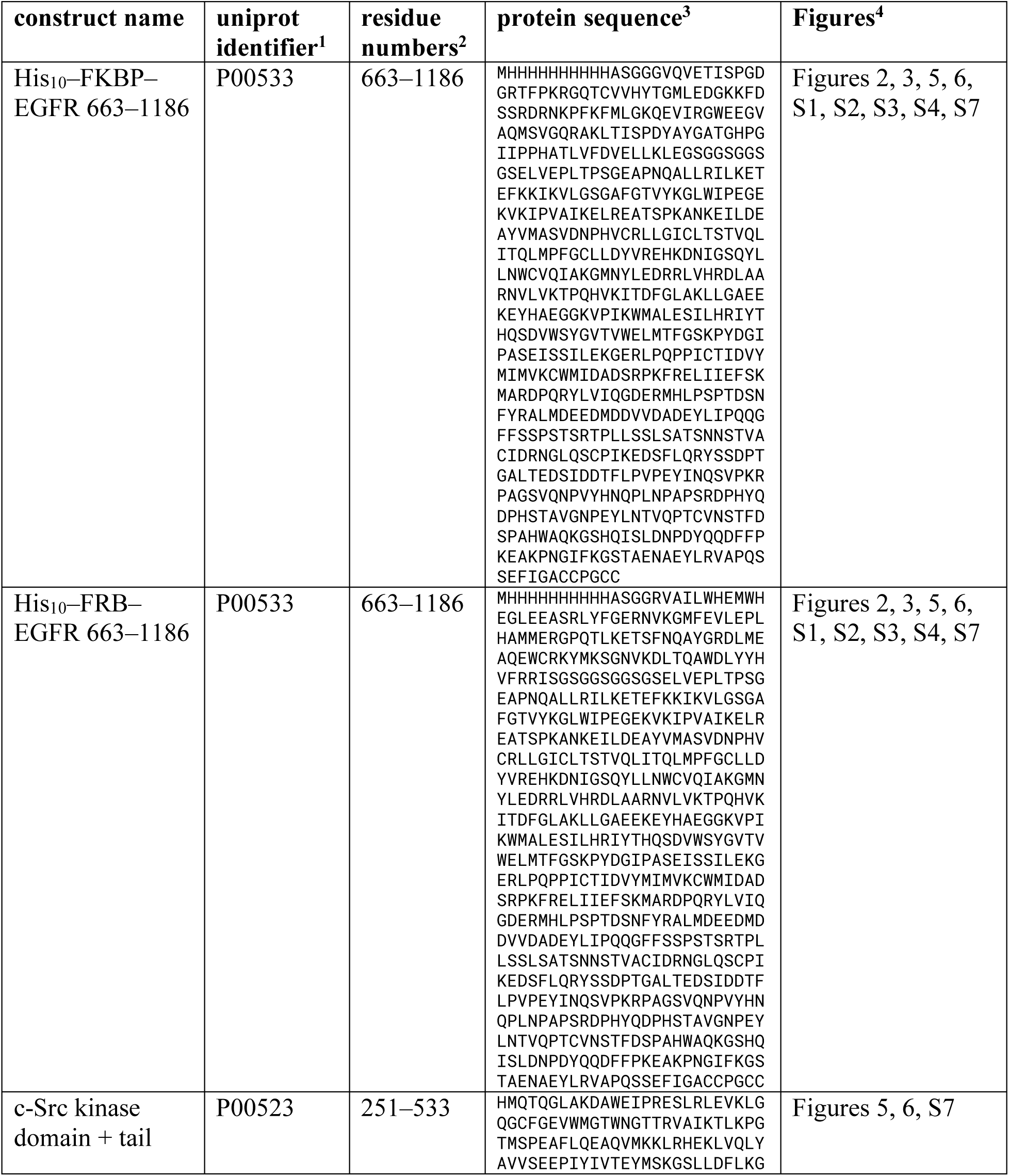
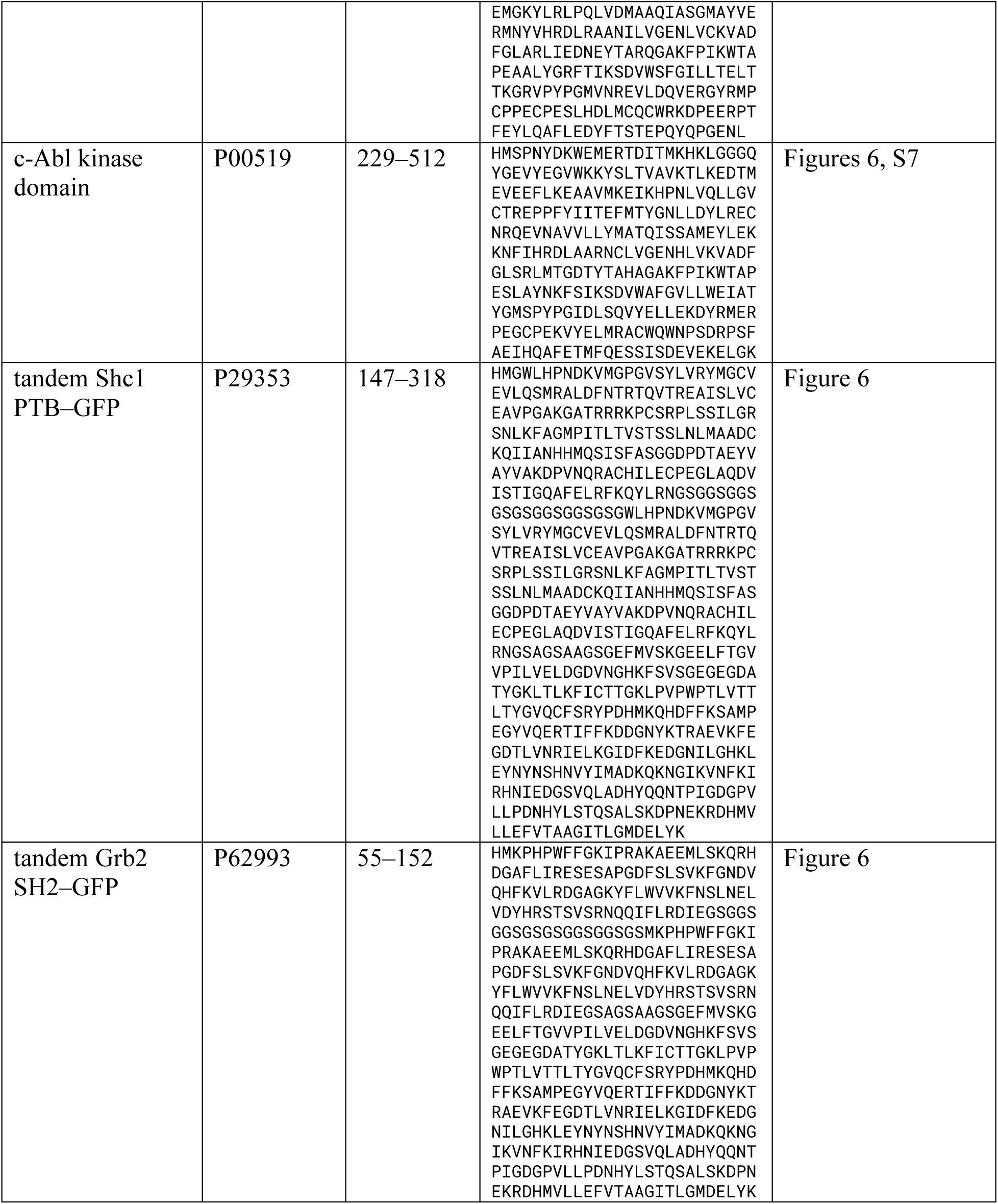
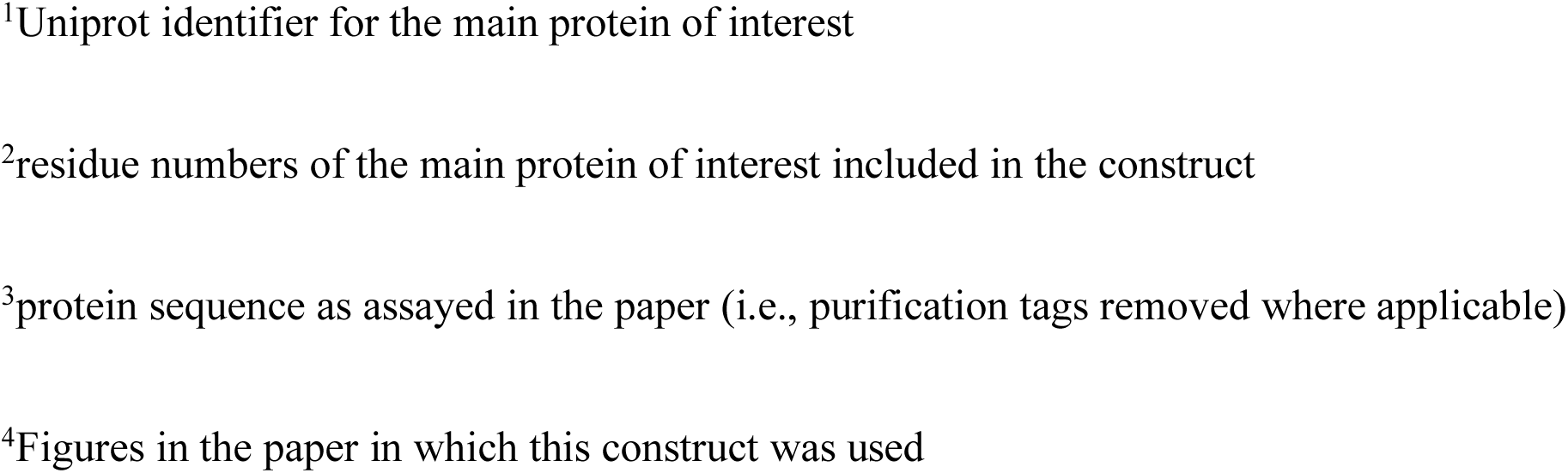
Sequences of purified proteins ^1^Uniprot identifier for the main protein of interest ^2^residue numbers of the main protein of interest included in the construct ^3^protein sequence as assayed in the paper (i.e., purification tags removed where applicable) ^4^Figures in the paper in which this construct was used

**Supplemental Table S2:**
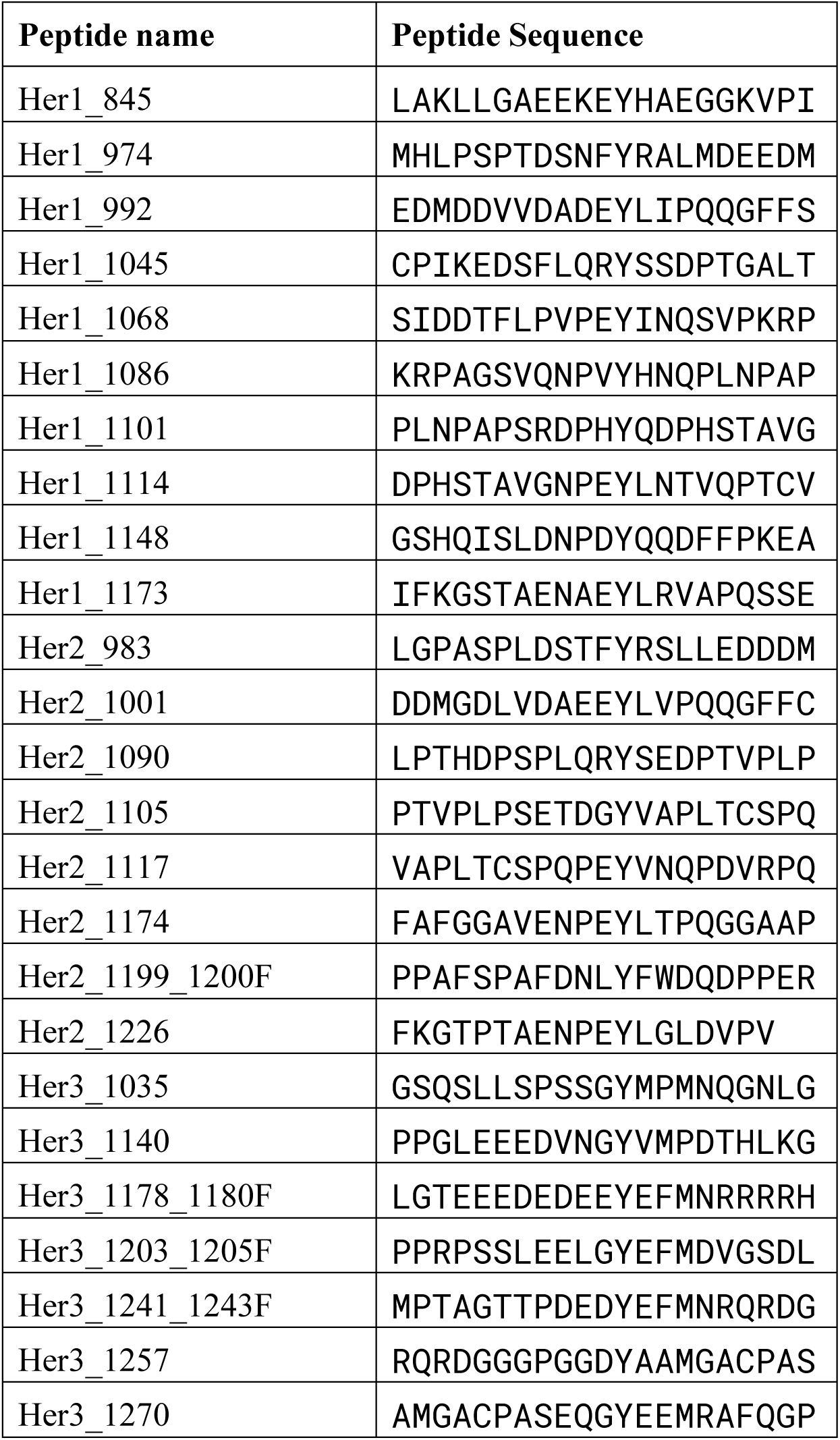
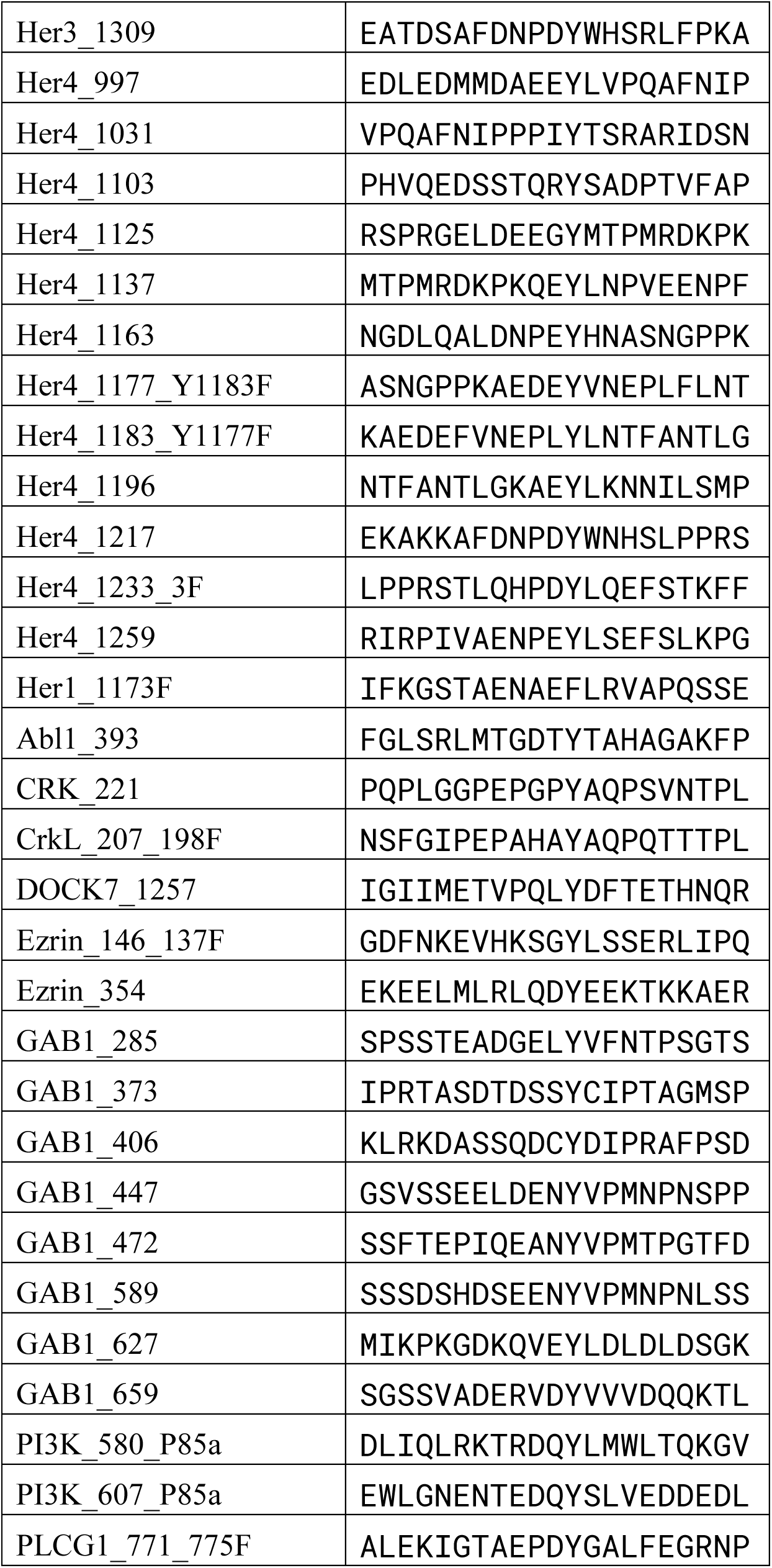
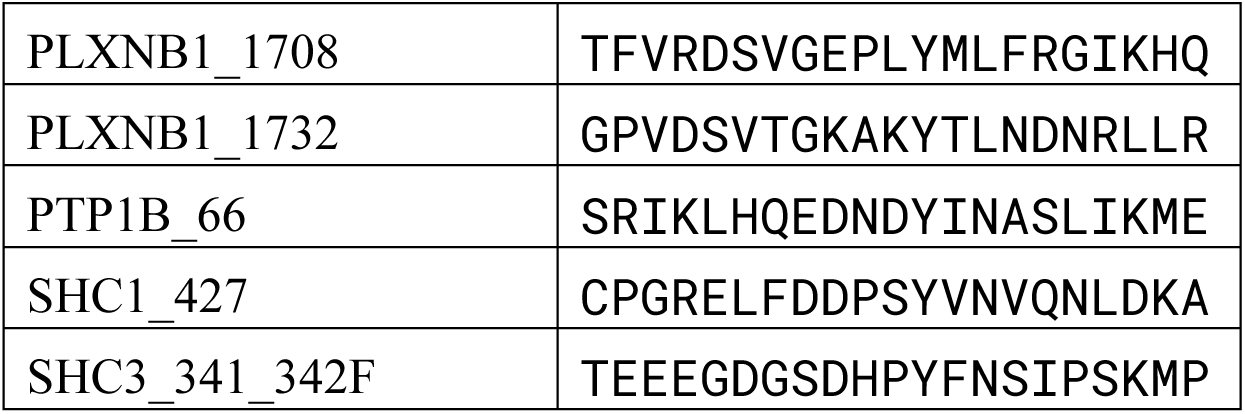
EGFR substrate phosphosite library used in Figure 5A and 5B

